# The Epithelial Na^+^ Channel UNC-8 promotes an endocytic mechanism that recycles presynaptic components from old to new boutons in remodeling neurons

**DOI:** 10.1101/2022.11.29.518248

**Authors:** Andrea Cuentas-Condori, Siqi Chen, Mia Krout, Kristin Gallick, John Tipps, Leah Flautt, Janet E. Richmond, David M. Miller

## Abstract

**Summary:** Presynaptic terminals are actively relocated during development to refine circuit function, but the underlying cell biological mechanisms are largely unknown. In *C. elegans*, the presynaptic boutons of GABAergic DD neurons are moved to new locations during early larval development. We show that developmentally regulated expression of a presynaptic Epithelial Na^+^ Channel (ENaC), UNC-8, promotes a Ca^2+^-dependent mechanism, resembling Activity-Dependent Bulk Endocytosis (ADBE), that dismantles presynaptic material for reassembly at nascent DD synapses. ADBE normally functions in highly active neurons to accelerate local recycling of synaptic vesicles. We show that DD presynaptic remodeling depends on canonical features of ADBE including elevated intracellular Ca^2+^, the phosphatase Calcineurin and its targets, dynamin and the F-BAR protein syndapin, and Arp2/3-driven actin polymerization. Thus, our findings suggest that a native mechanism (ADBE) for maintaining neurotransmitter release at local synapses has been repurposed, in this case, to dismantle presynaptic terminals for reassembly at new locations.

**Highlights:**

- Developing GABAergic neurons dismantle presynaptic terminals for reassembly at new locations.
- The DEG/ENaC protein, UNC-8, promotes presynaptic disassembly and recycling
- Ca^2+-^dependent endocytosis drives presynaptic disassembly and recycling to new boutons

## Introduction

Neural circuits are actively refined during development with the assembly of new synapses as others are removed. In the mammalian visual circuit, for example, axons of retinal ganglion cells (RGCs) initially project to the thalamus to innervate multiple targets with widely dispersed presynaptic boutons. In an activity-dependent mechanism that correlates with developmental age, distal boutons are eliminated while proximal RGC presynaptic domains are enlarged. Notably, the denuded sections of RGC axons are retracted at a later stage after RGC boutons have been relocated. Thus, synaptic refinement in this visual circuit is temporally segregated from axonal pruning and therefore likely depends on a separate mechanism (Hong et al., 2014).

Presynaptic boutons in the adult nervous system are also subject to remodeling and this inherent plasticity has been proposed to mediate recovery from injury (Berry & Nedivi, 2016; Holtmaat & Svoboda, 2009; Tennant et al., 2017). Live-imaging of mammalian neurons revealed that presynaptic varicosities are highly dynamic under basal conditions and also responsive to activity (Carrillo et al., 2013; De Paola et al., 2006). Thus, a better understanding of the mechanisms that govern presynaptic elimination and formation should advance the overarching goal of manipulating presynaptic structural plasticity to restore circuits damaged by injury or neurodegenerative disease.

Because synaptic remodeling is observed across species (Cuentas-Condori & Miller, 2020) and therefore likely to depend on conserved pathways, we are investigating the mechanism in the model organism, *C. elegans*, in which we can exploit powerful genetic strategies and live imaging methods. In the newly hatched larva (L1), DD class GABAergic motor neurons innervate ventral muscles. During the transition from the L1 to L2 larval stages, ventral DD synapses are eliminated, and new DD inputs are established with dorsal muscles (Cuentas-Condori & Miller, 2020; Hallam & Jin, 1998; Kurup & Jin, 2016; White et al., 1978) (Figure 1A). The relocation of DD presynaptic boutons proceeds without any evident alterations in external DD morphology, pointing to an internal cell biological mechanism that drives synaptic reorganization but not axonal plasticity (Mulcahy et al., 2022; White et al., 1978).

**Figure 1:**
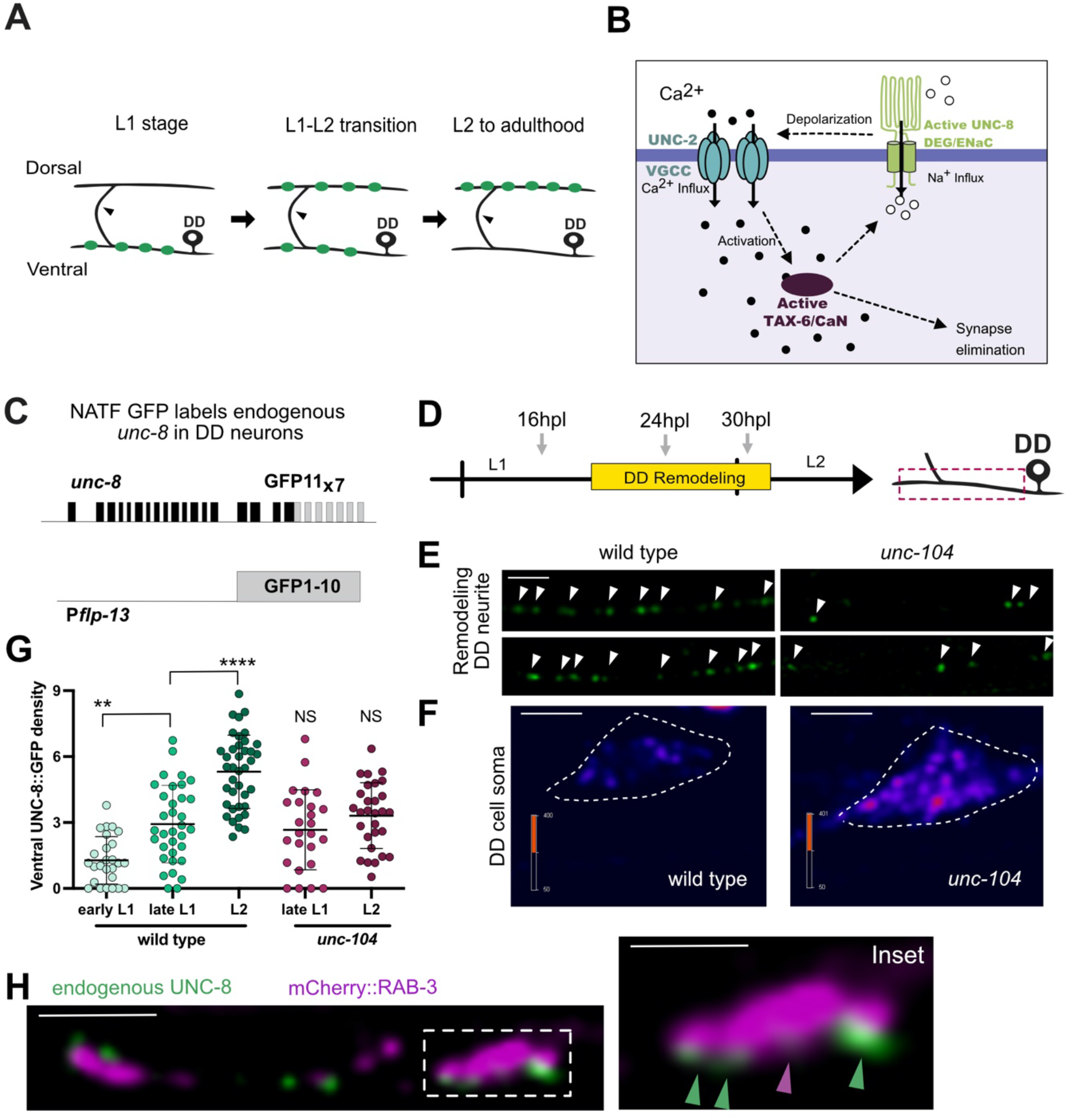
UNC-8 localizes to perisynaptic regions of remodeling DD presynaptic boutons. **A**. Re-location of presynaptic boutons (green) from ventral to dorsal DD neurites during early larval development (late L1 and L2 stages). Arrowheads denote the DD commissure. **B**. Working model depicts a positive feedback loop involving DEG/ENaC/UNC-8 channel, a Voltage-Gated Ca++ Channel (VGCC) and Calcineurin (TAX-6/CaN) that upregulates presynaptic Ca^2+^ for elimination of DD presynaptic terminals (Miller-Fleming et al., 2016). **C**. Split GFP strategy (NATF) (He et al., 2019) to tag endogenous UNC-8 in DD neurons. Seven copies of GFP11 (GFP11_x7_) were inserted at the endogenous *unc-8* locus to produce a C-terminal UNC-8::GFP11_x7_ fusion protein. An extrachromosomal array was used to drive the complementary GFP1-10 peptide in DD neurons under the *flp-13* promoter (P*flp-13*). **D**. Endogenous UNC-8::GFP puncta were evaluated (left) at developmental time points (arrows) spanning the remodeling period: early L1 (16hpl), late L1 (24 hpl) and L2 stages (32 hpl) (right) in remodeling DD axons. hpl = hours post lay (see Methods). **E**. Representative images of UNC-8::GFP puncta at the L2 stage in the wild type (left) vs *unc-104*/Kif1A mutant (right). Scale bar = 2 μm. Fix heatmap scales. **F**. Quantification of UNC-8::GFP density (UNC-8 puncta/10μm) at each timepoint shows progressive upregulation in the wild type (left): early L1 (1.28 ± 1.1, n=26), late L1 (2.93 ± 1.8, n=34), L2 stages (5.31 ± 1.7, n=39) but not in an *unc-104*/Kif1A mutant (right): late L1 (2.66 ± 1.8, n=25) and L2 (3.31 ± 1.5, n=20). Data are mean ± SD. Kruskal-Wallis test. ** p < 0.01, **** p < 0.0001 and NS is Not Significant. All comparisons are relative to the 24hpl time-point in the wild type. **G**. Representative images showing the accumulation of UNC-8::GFP puncta in the DD cell soma of trafficking defective *unc-104*/Kif1A mutant. Color key denotes relative UNC-8::GFP signal intensity. Scale bar = 2 μm. **H**. (Left) Dual-color imaging during the remodeling period (late L1) detects endogenous UNC-8::GFP puncta (green) and neighboring ventral presynaptic DD boutons labeled with mCherry::RAB-3 region (magenta). Scale bar = 2 μm. (Right) Inset shows close-up of UNC-8::GFP puncta (green) adjacent to presynaptic region (magenta). Scale bar = 1 μm.

Importantly, DD presynaptic remodeling is transcriptionally regulated (Howell et al., 2015; J. Meng et al., 2017; Miller-Fleming et al., 2021; Petersen et al., 2011; Thompson-Peer et al., 2012) and accelerated by synaptic function (Miller-Fleming et al., 2016; Thompson-Peer et al., 2012) suggesting that the underlying mechanism depends on the intersection of a genetic program with circuit activity. This idea is consistent with the finding that the homeodomain transcription factor IRX-1/Iroquois drives expression of UNC-8, a member of the degenerin/epithelial sodium channel (DEG/ENaC) family, that promotes disassembly of ventral DD presynaptic domains (Miller-Fleming et al., 2016, 2021; Petersen et al., 2011). A reconstituted UNC-8 channel preferentially gates Na^+^ in *Xenopus* oocytes (Wang et al., 2013) which suggests that the native UNC-8/DEG/ENaC channel could depolarize the GABA neurons in which it is expressed. In turn, this effect is predicted to enhance Ca^2+^ import by presynaptic voltage gated calcium channels (VGCCs), an idea supported by the additional result that UNC-2/VGCC is required for UNC-8-dependent presynaptic disassembly (Miller-Fleming et al., 2016) (Figure 1B). This model parallels the finding that *Drosophila* Pickpocket/DEG/ENaC channel is presynaptically localized in motor neurons in a homeostatic mechanism that elevates intracellular Ca^2+^ for neurotransmitter release (Orr et al., 2017; Younger et al., 2013). The radically different outcome of synaptic destruction that arises from UNC-8 function in *C. elegans* depends on the serine/threonine phosphatase TAX-6/CalcineurinA and its regulatory subunit CNB-1/CalcineurinB. Calcineurin/CaN is activated by intracellular Ca^2+^ and genetic results suggest that it functions upstream of UNC-8. Thus, UNC-8/DEG/ENaC, UNC-2/VGCC and TAX-6/CaN may constitute a positive feedback loop to amplify Ca^2+^ influx to drive synaptic disassembly (Figure 1B)(Miller-Fleming et al., 2016). Because CaN is known to function as a key regulator of Activity-Dependent Bulk Endocytosis (ADBE) (Emma L Clayton & Cousin, 2009), we have investigated the possibility that an ADBE-like mechanism is involved in DD synapse elimination.

Synaptic vesicle (SV) membrane and proteins are recycled at presynaptic domains by parallel-acting pathways (Chanaday et al., 2019). In the canonical mechanism, single SVs are retrieved from the presynaptic membrane as clathrin-coated vesicles. In highly active neurons in which the plasma membrane is rapidly expanded by fusion with multiple SVs, recycling is also achieved by ADBE in which large (80-100 nm) endosomes recover SV membrane in a clathrin-independent mechanism (Chanaday et al., 2019; Emma L Clayton & Cousin, 2009). ADBE depends on CaN, which induces assembly of the Dynamin-Syndapin complex (Anggono et al., 2006; E. L. Clayton et al., 2009; Emma L Clayton & Cousin, 2009; Cousin & Robinson, 2001). The initial formation of bulk endosomes requires actin (Wu et al., 2016) and has been proposed to depend on branched actin polymerization triggered by Syndapin recruitment of N-WASP to the remodeling membrane (Kessels & Qualmann, 2002, 2004). N-WASP is a well-known activator of the branched-actin nucleator Arp2/3 (Rohatgi et al., 1999). Consistent with this finding, Syndapin and Arp2/3 subunits are enriched in bulk endosomes derived from highly active neurons (A C Kokotos et al., 2018). Other studies in neurosecretory cells determined that the maturation of bulk endosomes depends on an acto-myosin II ring that separates the invaginating bulk endosomes from the membrane (Gormal et al., 2015). Thus, the actin cytoskeleton may participate in both the formation and scission of bulk endosomes.

Here we show that UNC-8/DEG/ENaC and TAX-6/CaN localize to the presynaptic regions of remodeling DD neurons where UNC-8 drives local Ca^2+^ transients that are strengthened by TAX-6 function, results consistent with the proposal that UNC-8 and TAX-6 act in a positive feedback loop to elevate presynaptic Ca^2+^(Miller-Fleming et al., 2016). In experiments to test the idea that an ADBE-like mechanism functions downstream of UNC-8, we show that canonical ADBE components including CaN, Dynamin, Syndapin and Arp2/3 are required for UNC-8-dependent removal of DD presynaptic components. Importantly, our results provide a possible explanation for the dramatically different outcomes of ADBE, which maintains synaptic function, versus our proposed bulk endocytic mechanism, which dismantles the synapse. As noted above, in ADBE, bulk endosomes give rise to synaptic vesicles that replenish the local pool. Our studies of remodeling DD neurons have revealed that ventrally located presynaptic components are also recycled, but in this case to distal synapses in the dorsal DD neurite. These findings are important because they suggest that a conserved endocytic mechanism could be widely utilized to couple synapse elimination with synapse formation during circuit refinement.

## Results

### Kinesin-3-dependent transport of the DEG/ENaC channel protein UNC-8 to remodeling presynaptic boutons

The transcript for the Epithelial Na channel UNC-8 is upregulated by the IRX-1/Iroquois transcription factor in remodeling DD GABAergic neurons where it promotes the elimination of presynaptic termini (Miller-Fleming et al., 2021; Petersen et al., 2011). We showed that UNC-8::GFP protein expressed from a transgenic array labels ventral DD neurites (Miller-Fleming et al., 2016), but this result did not establish the timing of endogenous UNC-8 expression in DD neurons and ventral localization of UNC-8::GFP could be misleading due to over-expression.

To monitor the endogenous UNC-8 protein during DD remodeling, we used NATF, a two-component system (GFP1-10 and GFP11) that exploits the self-annealing property of complementary split-GFP peptides (He et al., 2019; Kamiyama et al., 2016). We attached seven copies of GFP11 to the *unc-8* locus to amplify the GFP signal and expressed the complementary GFP1-10 peptide in DD neurons (Figure 1C). We used this strain to visualize expression of the endogenous UNC-8 protein at three different time-points, before remodeling (16 hpl), early remodeling (24 hpl) and late remodeling (30 hpl) (Figure 1D, Figure S1). Dim UNC-8::GFP puncta were sparsely detectable in DD neurons in early L1 larvae (16 hpl) but were robustly elevated as DD remodeling progressed in the late L1 (24 hpl) and early L2 larval stages (30 hpl) (Figure 1E). This finding is consistent with the idea that *unc-8* is transcriptionally upregulated to elevate UNC-8 protein for DD remodeling(Miller-Fleming et al., 2021).

To investigate how UNC-8 is transported in remodeling DD axons, we evaluated the distribution of UNC-8::GFP puncta in a mutant that disrupts cargo-loading function of the kinesin-3 protein UNC-104/KIF1A (Kumar et al., 2010). In *unc-104* mutants, synaptic vesicles fail to populate the axon and accumulate in the cell soma (Hall & Hedgecock, 1991; Otsuka et al., 1991). Strikingly, UNC-8::GFP puncta are sequestered in DD cell soma in UNC-104/KIF1A mutant animals (Figure 1F-G). These results demonstrate that robust trafficking of UNC-8 protein into remodeling DD axons requires UNC-104/KIF1A activity.

Because UNC-104/KIF1A transports synaptic vesicle components from the cell soma to the presynaptic apparatus, we reasoned that UNC-8 might also localize to DD presynaptic domains. To test this idea, we used a two-color imaging strategy to compare the localization of UNC-8::GFP vs RAB-3 labeled with mCherry. During the DD remodeling period (24 hpl), endogenous UNC-8::GFP puncta appeared in regions adjacent to mCherry::RAB-3 clusters in the remodeling axon (Figure 1H). A similar result was obtained for members of the *ppk* DEG/ENaC family in *Drosophila* which also localize to perisynaptic regions of glutamatergic motor neuron boutons (Orr et al., 2017). Together, our results demonstrate that UNC-8 is developmentally upregulated and localizes to the vicinity of remodeling DD boutons in a kinesin-3-dependent manner. Interestingly, UNC-8::GFP remains in the ventral DD neurite (Figure S1D) after presynaptic boutons are relocated to the dorsal side, a result that could be important for maintaining the stability of dorsal DD synapses in the adult.

### UNC-8 maintains Ca^2+^ levels in remodeling DD neurons

Previous work established that a reconstituted UNC-8 channel mediates Na^+^ influx (Wang et al., 2013). The resultant depolarization of local membrane potential in DD presynaptic boutons is predicted to enhance VGCC/UNC-2 channel activation and consequently elevate intracellular Ca^2+^ (Figure 1B) (Miller-Fleming et al., 2016). To test this idea *in vivo*, we expressed the Ca^2+^ sensor, GCaMP6s, in DD neurons to track spontaneous Ca^2+^ changes at remodeling synapses (26 hpl) (Figure 2A). In wild-type animals, we detected striking Ca^2+^ fluctuations confined to bouton-like structures (Figure 2B) that likely correspond to DD presynaptic terminals (Kittelmann et al., 2013) (Video S1). Live imaging revealed that at basal states, GCaMP fluorescence is downregulated in *unc-8* mutants (Figure 2C-D), suggesting that UNC-8 maintains Ca^2+^ levels in remodeling DD neurons (Video S2).

**Figure 2:**
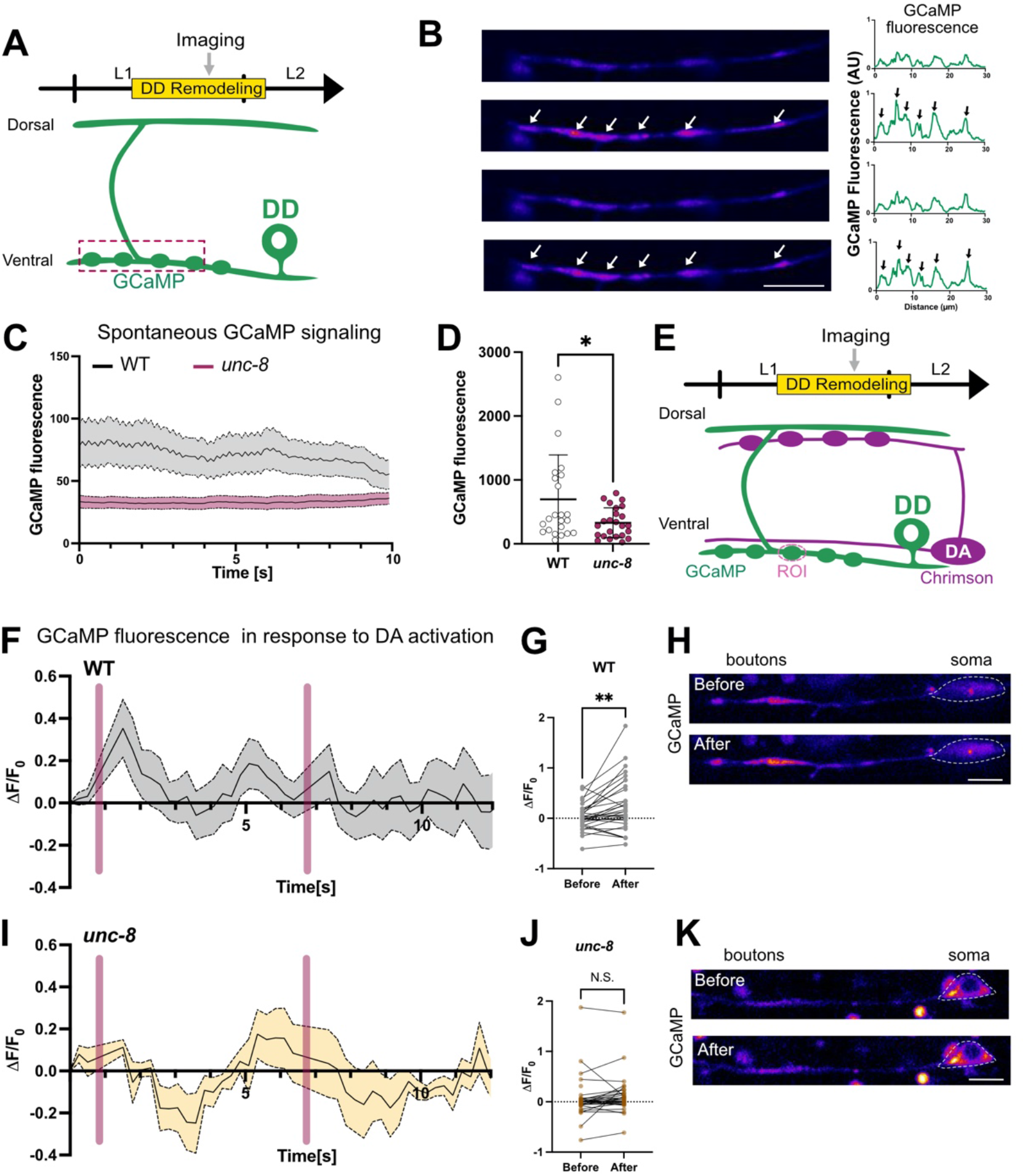
UNC-8/ENaC promotes Ca^++^ transients in presynaptic boutons during DD remodeling. **A**. Experimental set-up utilizes GCaMP (green) to monitor Ca^++^ transients in DD neurons. Imaging was performed on synchronized L1 larvae during the remodeling window (26-28 hpl) (see Figure 1B-C). **B**. (Left) Snapshots of spontaneous GCaMP signal in a wild-type remodeling ventral DD neurite with sequential images shown at 1 fps (frame per second) and (Right) corresponding line scans. Arrows point to bouton-like regions of elevated Ca^++^ signal. Scale bar = 5 μm. **C-D**. Spontaneous Ca^++^ fluctuations were detected in wild-type (WT) and *unc-8* mutant animals. GCaMP fluorescence is elevated in WT (695 ± 145, n=23) vs *unc-8* (330 ± 47.7, n=24). Mann-Whitney test, * p = 0.0343. **E**. Experimental set-up for measuring evoked Ca^++^ transients in remodeling ventral DD neurites: (Top) Cholinergic DA motor neuron presynaptic boutons (magenta) provide excitatory input to DD GABAergic motor neurons (green) in the dorsal nerve cord. The red-light activated channelrhodopsin, Chrimson, was expressed in DA neurons (magenta) to evoke Ca^++^ transients in postsynaptic DD neurons (Panels D-H). GCaMP signals were collected from individual DD presynaptic boutons (ROI). **F-H. (F)** Chrimson activation in DA neurons with a pulse of 561nm light (magenta bars) evokes GCaMP transients in wild-type (WT) DD presynaptic boutons. **(G)** GCaMP signal before pulse of 561 nm light (0.02 ± 0.3, n=32) is elevated (0.25 ± 0.5, n=32) after Chrimson activation. Non-parametric paired Wilcoxon test ** p = 0.002. **(H)** Snapshots of a remodeling bouton (ROI) shows that GCaMP levels are upregulated after Chrimson activation in the wild type. Dashed circle denotes DD cell soma. Scale bar = 4 μm. **I-K. (I)** Chrimson activation of *unc-8* mutant DA neurons fails to evoke a GCaMP response in DD neurons. **(J)** GCaMP baseline (0.06 ± 0.4, n=32) does not increase after DA activation (0.07 ± 0.5, n=32). Non-parametric paired Wilcoxon test, p = 0.0803. **(K)** Snapshots of a remodeling DD bouton (ROI) show that GCaMP levels do not increase upon Chrimson activation in *unc-8* mutants. Dashed circle denotes DD cell soma. Scale bar = 4 μm. All animals were grown in the presence of ATR.

To determine if UNC-8 can also mediate evoked synaptic responses, we activated presynaptic cholinergic DA motor neurons with the light-sensitive opsin, Chrimson (Schild & Glauser, 2015), and monitored Ca^2+^ changes in DD neurons with GCaMP6s (Figure 2E). This experiment detected consistent upregulation of Ca^2+^ at remodeling DD boutons with DA neuron activation in wild-type animals (Figure 2F-H). Importantly, this effect depends on ATR, a necessary channelrhodopsin cofactor (Figure S2A-B). In *unc-8* mutants, however, Ca^2+^ levels in DD neurons were not elevated in response to DA activation either in the presence (Figure 2I-K) or absence of ATR. (Figure S2C-D) (Video S3). This result parallels a previous finding in which evoked Ca^2+^ responses are reduced in presynaptic regions by the ENaC channel antagonist, Benzamil, presumptively due to inhibition of a presynaptic PPK11/16 containing DEG/ENaC channel at the *Drosophila* neuromuscular junction (Younger et al., 2013). Thus, our results lend strong support to the proposal that the presynaptic UNC-8 channel mediates elevated Ca^2+^ levels in remodeling boutons and substantiates a role for synaptic activity, through Ca^2+^ signaling, in disassembly of DD presynaptic domains (Figure 1B) (Miller-Fleming et al., 2016).

### Calcineurin/CaN localizes to presynaptic terminals in DD neurons

We have previously shown that the conserved phosphatase Calcineurin/CaN functions with *unc-8* in a common genetic pathway to promote presynaptic disassembly in remodeling DD neurons (Miller-Fleming et al., 2016). Because we detected UNC-8 protein at remodeling DD presynaptic boutons (Figure 1F), we devised an experiment to determine if TAX-6, the catalytic subunit of CaN in *C. elegans*, is presynaptically localized. To visualize endogenous TAX-6, we used the NATF strategy (He et al., 2019) to insert 3 copies of GFP11 at the *tax-6* locus and expressed the complementary GFP1-10 peptide in DD neurons (Figure 3A). Imaging before the remodeling period (L1 larval stage) revealed that endogenous TAX-6 is enriched in the DD ventral neurite. In contrast, after remodeling, at the L4 stage, TAX-6::GFP is brightest on the dorsal side (Figure 3B-C). This localization pattern for TAX-6::GFP matches that of DD presynaptic domains during larval development (Figure 3D-E). Close inspection revealed both a diffuse TAX-6::GFP signal as well as locally enriched TAX-6::GFP puncta throughout the axon (Figure 3D-E). To determine if TAX-6::GFP puncta are localized to presynaptic terminals, we labeled the active zone protein, CLA-1, with mRuby (Xuan et al., 2017). Imaging revealed that TAX-6::GFP puncta are consistently enriched at CLA-1::mRuby-labeled active zones (Figure 3D-E). Thus, our results suggest that both UNC-8 and TAX-6 are localized to presynaptic regions in remodeling DD neurons. As noted above, however, UNC-8 is strictly localized to perisynaptic regions on the ventral side throughout development and in adults (data not shown) whereas CaN is relocated to dorsal DD boutons (Figure 3E).

**Figure 3:**
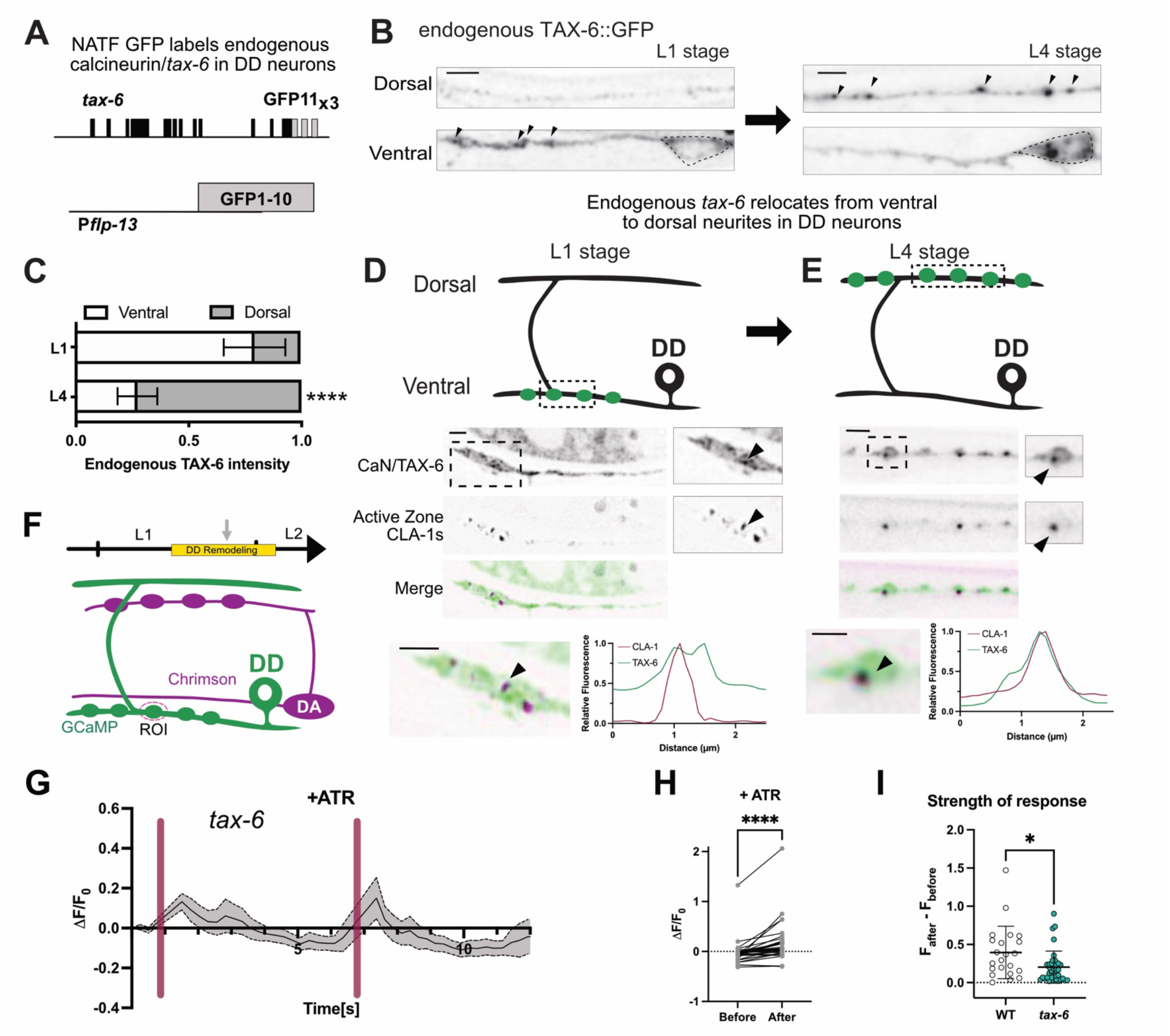
Presynaptic TAX-6/CaN elevates Ca^++^ transients in remodeling DD boutons. **A**. Split GFP (NATF) strategy for labeling endogenous CaN/TAX-6 in DD neurons. Three copies of GFP11 were inserted at the endogenous *tax-6* locus to produce a C-terminal TAX-6:: GFP11_x3_ fusion protein. An extrachromosomal array was used to drive the complementary GFP1-10 peptide in DD neurons under the *flp-13* promoter (*Pflp-13*). **B-C. (B)** (Left) Before DD remodeling, at the L1 stage, endogenous GFP-labeled TAX-6 is enriched (arrowheads) in the ventral DD neurite but is relocated to the dorsal cord (right) after remodeling, at the L4 stage. Scale bar = 2 μm. **(C)** Quantification of TAX-6::GFP detects initial enrichment in the ventral cord (V = 0.79 ± 0.1 vs D = 0.2 ± 0.1, n=29) in L1 larvae and later enrichment on the dorsal side at the L4 stage (V = 0.27 ± 0.1, n=21 vs D = 0.73 ± 0.1, n-21). One-Way ANOVA with Tukey’s multiple comparisons test, **** p < 0.0001. **D-E** (Top) Graphical representation of the relocation of TAX-6 from ventral (L1 stage) to dorsal (L4 stage) DD neurites. (Bottom) CaN/TAX-6 is enriched at presynaptic active zones. Co-localization (arrowheads) of endogenous CaN/TAX-6::GFP (green) and active zone protein CLA-1s (magenta) **(D)** on the ventral side before remodeling (L1 stage) and **(E)** on the dorsal side (L4 stage) after remodeling. (bottom) Line scans of insets show co-localization of CaN/TAX-6 (green) and CLA-1s (magenta). Scale bar = 2 μm. **F**. Experimental set-up: (Top) The red-light activated channelrhodopsin, Chrimson, was expressed in presynaptic DA neurons (magenta) for experiments that evoke Ca^++^ transients in DD neurons (Panels D-H). **G-H. (G)** In the presence of ATR (+ATR), Chrimson activation in DA neurons with a pulse of 561nm light (magenta bar) elevates GCaMP fluorescence in remodeling DD presynaptic boutons (ROI) (H) (before activation = −0.02 ± 0.2, n=44, after activation = 0.14 ± 0.4, n=44). Non-parametric paired Wilcoxon test, **** p < 0.0001. **(I)** GCaMP fluorescence is elevated (normalized to F_0_) after Chrimson activation in wild-type (WT) (0.04 ± 0.3, n=44) in comparison to *tax-6* mutants (0.04 ± 0.3, n=44). Mann-Whitney test, * is p = 0.0109.

### Calcineurin/CaN upregulates Ca^2+^ in remodeling boutons

Because we found that UNC-8 maintains both intrinsic and evoked Ca^*2+*^ responses in remodeling DD neurons (Figure 2), we predicted that TAX-6 might also regulate intracellular Ca^2+^. To test this idea, we used GCaMP to monitor Ca^2+^ levels in remodeling DD boutons in *tax-6* mutants. In contrast to *unc-8* mutants in which Ca^2+^ transients are largely abrogated (Figure 2I-J), intrinsic Ca^2+^ levels in *tax-6* mutants are comparable to that of wild type (Figure S3A-B). To evaluate the role of TAX-6 in evoked synaptic function, we used Chrimson to activate presynaptic DA neurons and monitored Ca^2+^ in DD neurons with GCaMP6s (Figure 3F). This experiment revealed that evoked Ca^2+^ transients were observed in *tax-6* mutants (Figure 3G-H) and that this effect is dependent on ATR (Figure S3C-D). However, the intensity of the evoked Ca^2+^ response (F_after_-F_before_) in *tax-6* mutant animals is significantly reduced in comparison to wild type (Figure 3I). These findings are consistent with the idea that TAX-6 strengthens the Ca^2+^ response, perhaps by tuning *unc-8* activity (Figure 1B) (Miller-Fleming et al., 2016).

### Calcineurin/CaN functions downstream of UNC-8 to trigger presynaptic disassembly

We have shown that TAX-6, the catalytic subunit of CaN, localizes to DD presynaptic termini (Figure 3A-E) and that it contributes to the evoked Ca^2+^ transients in remodeling DD neurons (Figure 3 F-I). Together, these findings are consistent with the idea that TAX-6/CaN functions in a positive feedback loop with ENaC/UNC-8 and VGCC/UNC-2 to elevate intracellular Ca^2+^ for disassembly of the DD presynaptic apparatus (Miller-Fleming et al., 2016) (Figure 1B). This model also predicts that a loss-of-function mutation in *tax-6/CaN* should impair ENaC/UNC-8-dependent presynaptic disassembly. In the wild type, Ventral D (VD) GABAergic neurons do not express ENaC/UNC-8 and do not undergo synaptic remodeling. Forced expression of ENaC/UNC-8 in VD neurons, however, is sufficient to drive the removal of the synaptic vesicle-associated protein Synaptobrevin/SNB-1::GFP from VD synapses (Figure 4B) (Miller-Fleming et al., 2016, 2021). We exploited this experimental paradigm to show that RNAi knock down of *tax-6/CaN* prevents UNC-8-dependent elimination of VD synapses (Figure 4C), a result consistent with the hypothesis that TAX-6/CaN functions with UNC-8 to dismantle the GABAergic presynaptic apparatus (Figure 1B).

**Figure. 4.**
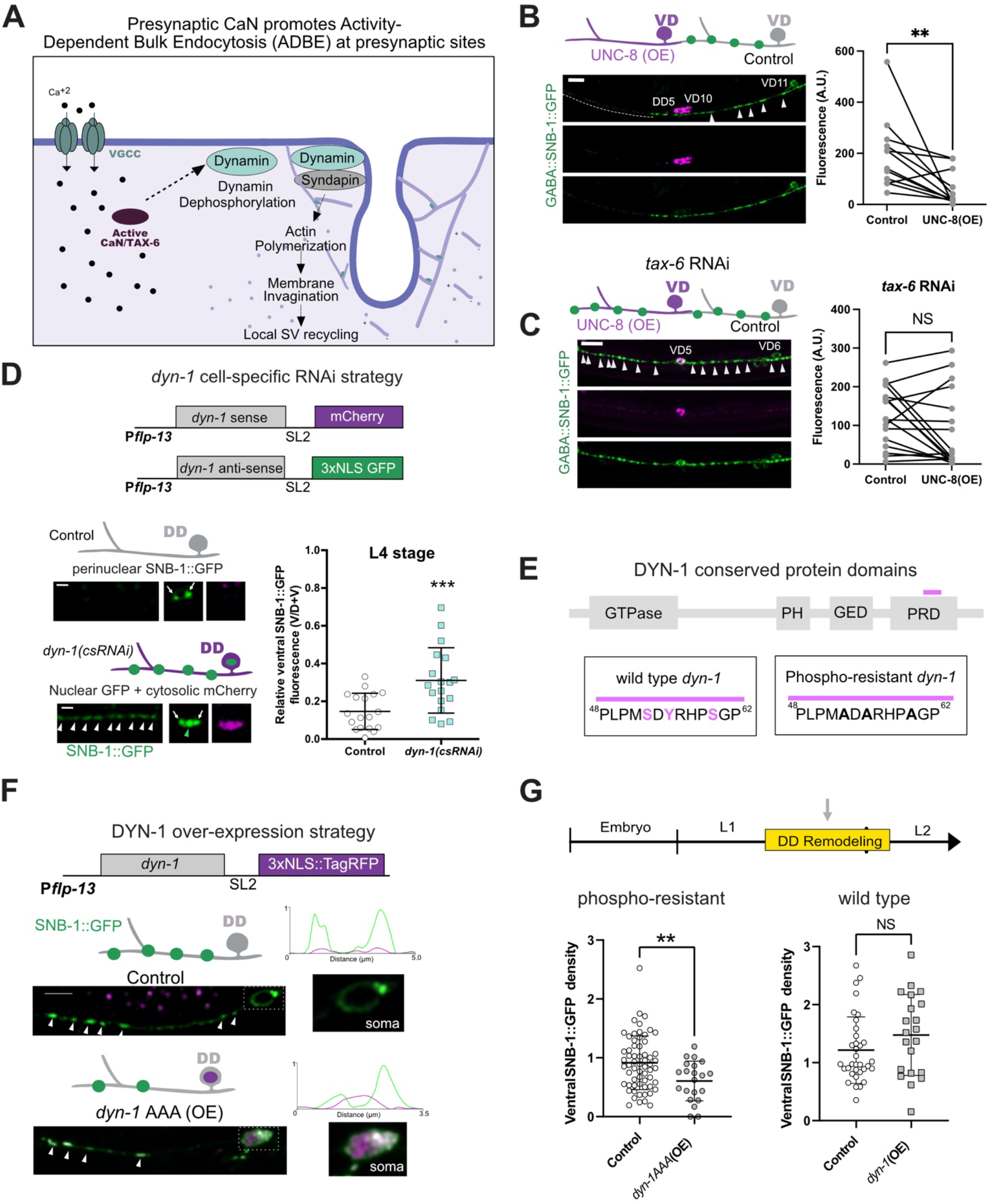
Cell autonomous dynamin activity is required for presynaptic disassembly in remodeling DD neurons. **A**. During Activity-Dependent Bulk Endocytosis (ADBE), intense synaptic activity elevates intracellular Ca^++^ to activate CaN/TAX-6. Dephosphorylation of dynamin by CaN promotes the formation of the dynamin-syndapin complex which localizes to the membrane to drive branched actin polymerization for bulk endosome formation and local synaptic vesicle (SV) recycling (Emma L Clayton & Cousin, 2009). **B**. (Left) Over-expression (OE) of UNC-8 in VD neurons induces presynaptic disassembly. Control VD neuron (VD11, gray) with prominent SNB-1::GFP puncta (arrowheads) vs anterior region of UNC-8(OE) VD neuron (V10, magenta) with fewer SNB-1::GFP puncta (dashed line). (Right) Paired analysis of neighboring VD neurons: Control VDs (196 ± 139 A.U, n=12) and UNC-8(OE) VD neurons (56.7 ± 67 A.U., n=12). Wilcoxon matched-pairs signed rank test, **p = 0.0024. **C**. TAX-6 is required for UNC-8-dependent removal of presynaptic domains in VD neurons. (Left) *tax-6* RNAi-treated UNC-8(OE) VD neuron (VD5, magenta) vs adjacent control VD neuron (VD6, gray) both with prominent SNB-1::GFP puncta (arrowheads). (Right) Paired analysis of neighboring VD neurons treated with *tax-6* RNAi: Control VDs (126 ± 76 A.U., n=17) and UNC-8 (OE) VD neurons (86 ± 99 A.U., n=17). Wilcoxon matched-pairs signed rank test, p = 0.16. NS is Not Significant. Scale bars = 10 μm. L4 larvae imaged for B and C. **D**. (Top) Transgenic strategy for cell-specific knockdown of *dyn-1(csRNAi)* (See Methods). (Middle, Left) Control DD neuron that does not express the *dyn-1(csRNAi)* transgenic array (grey) shows perinuclear SNB-1::GFP (arrows). (Bottom left) *dyn-1(csRNAi*)-expressing DD neurons (magenta) are labeled with cytosolic mCherry, nuclear GFP (green arrowhead), perinuclear SNB-1::GFP (arrows) and retain SNB-1::GFP puncta (white arrowheads) in the ventral nerve cord. (Bottom, Right) Fraction of SNB-1::GFP fluorescence in ventral DD neurites (V/V+D = Ventral/Ventral + Dorsal) showing that *dyn-1(csRNA)-*treated DD neurons retain a greater fraction of ventral SNB-1::GFP fluorescence (0.31 ± 0.2, n=18) than controls (0.15 ± 0.1, n=18). Unpaired t-test, *** is p = 0.0006. Scale bar = 10 μm. **E**. Schematic of conserved DYN-1 protein domains (see text for details). Magenta bar denotes location of C-terminal putative dynamin phospho-box sequence (Zielinska et al., 2009) with phosphorylatable residues (S53, Y55, S59) in wild type and converted to Alanine (A) in phospho-resistant *dyn-1* mutant. **F**. (Top) Transgenic strategy for over-expression of Dynamin (DYN-1) and phospho-resistant DYN-1 in DD neurons (See Methods). (Middle) Control DD neuron (grey) that does not express DYN-1 shows robust SNB-1::GFP puncta (white arrowheads). Inset of cell soma and line tracing. (Bottom) DD neuron that overexpresses phospho-resistant DYN-1 [*dyn-1AAA (OE)*] is labeled with nuclear-localized TagRFP (magenta) (Inset) and shows fewer SNB-1::GFP puncta than control DD neurons (white arrowheads). Scale bar = 10 μm, late L1 larval stage. **G**. (Top) L1 larvae were imaged during the remodeling window (arrow) to assess ventral SNB-1::GFP in DD neurons. (Bottom-Left) Over-expression of phospho-resistant DYN-1 [*dyn-1AAA(OE)*] accelerates SNB-1::GFP removal (0.61 ± 0.3, n=21) vs control DD neurons (0.92 ± 0.5, n=62). (Bottom Right) Over-expression of wild-type dynamin [dyn-1(OE)} (1.47 ± 0.7, n=21) does not result in fewer SNB-1::GFP puncta vs control DD neurons (1.21 ± 0.6, n=34). Density = SNB-1 puncta/10 micron.

### Dephosphins, targets of CaN phosphatase activity, drive the removal of presynaptic components in remodeling DD neurons

In addition to amplifying Ca^2+^ import, potentially by enhancing UNC-8 channel activity, CaN/TAX-6 could also regulate additional downstream effectors for presynaptic disassembly. One of the known roles for CaN at presynaptic termini is to trigger Activity-Dependent Bulk Endocytosis (ADBE) during periods of high synaptic activity (Figure 4A). In this mechanism, elevated calcium activates CaN to function as a Ca^2+^-dependent phosphatase for dephosphorylation of a well-characterized group of synaptic proteins known as dephosphins (Emma L Clayton & Cousin, 2009; Cousin & Robinson, 2001). For example, CaN dephosphorylates residues in a conserved phospho-box near the C terminus of the GTPase, Dynamin. The dephosphorylation of dynamin promotes the formation of the Dynamin-Syndapin complex (Anggono et al., 2006; Xue et al., 2011). Through its SH3 domain, Syndapin recruits components that drive branched-actin polymerization for membrane invagination to form presynaptic bulk endosomes (Kessels & Qualmann, 2002). Importantly, the endocytic mechanism in ADBE does not require clathrin (Emma L Clayton & Cousin, 2009), which is necessary for synaptic vesicle endocytosis during mild stimulation (Chanaday et al., 2019). The canonical role of ADBE is to recycle large amounts of membrane during periods of intense activity for the local replenishment of synaptic vesicles for neurotransmission (Emma L Clayton & Cousin, 2009). Here, we test the idea that TAX-6/CaN drives an ADBE-like mechanism that orchestrates the removal of presynaptic components from remodeling DD neurons.

During ADBE, CaN targets dephosphins to drive bulk endocytosis of the presynaptic membrane (Emma L Clayton et al., 2007). If an ADBE-like mechanism is activated by CaN/TAX-6 in remodeling GABAergic neurons, then dephosphins with known roles in ADBE should be necessary for synapse elimination. To test this prediction, we asked if conserved dephosphins (e.g., Amphiphysin/*amph-1*, Epsin/*epn-1*, Eps15/*ehs-1*, Synaptojanin/*unc-26i* and AP180/*unc-11*) are required for the removal of the presynaptic domain. For this experiment, we exploited the remodeling phenotype of *unc-55* mutants in which VD ventral synapses are ectopically eliminated. Normally the UNC-55 COUP-TF transcription factor functions in VD neurons to block the canonical DD remodeling program. Consequently, in wild-type adults, VD synapses are located in the ventral cord and DD GABAergic synapses are positioned on the dorsal side. In *unc-55* mutants, however, both DD and VD neurons remodel, thus resulting in the elimination of all GABAergic synapses from ventral muscles (Figure S4A-B) (Mimi Zhou & Walthall, 1998; Walthall et al., 1993; B. Yu et al., 2017). If a gene is necessary for disassembly of GABA synapses, then RNAi knockdown should suppress the Unc-55 phenotype by impeding the elimination of ventral synapses (Figure S4C)(Petersen et al., 2011; Thompson-Peer et al., 2012). In this case, we assessed the number of GABAergic synapses in the ventral nerve cord labeled with either Synaptobrevin/SNB-1::GFP or the active zone protein liprin-alpha/SYD-2::GFP. This candidate screen determined that Amphiphysin/*amph-1*, Epsin/*epn-1* and Eps15/*ehs-1* normally promote GABAergic presynaptic elimination (Figure S4E). A genetic mutant of the dephosphin, Synaptojanin/*unc-26*, also suppressed the Unc-55 remodeling phenotype (Figure S4E). Notably, the CaN target that drives clathrin-mediated endocytosis, AP180/*unc-11*, and the clathrin-related proteins, the clathrin light chain/*clic-1* and the clathrin adaptor AP2/*dpy-23*, are dispensable for the removal of remodeling GABAergic synapses (Figure S4D). Thus, our findings support the hypothesis that conserved CaN targets, (e.g, dephosphins) are required for a clathrin-independent endocytic mechanism that dismantles the presynaptic apparatus of remodeling GABAergic neurons.

### Phospho-resistant dynamin functions in DD neurons to promote presynaptic disassembly

The GTPase, Dynamin, is a known target of CaN and functions as a key effector of both ADBE (E. L. Clayton et al., 2009) and the canonical clathrin-dependent mechanism for recycling synaptic vesicle membrane at active synapses (Chanaday et al., 2019). To circumvent the lethal phenotype of Dynamin/*dyn-1* mutants in *C. elegans* (X. Yu et al., 2006), we devised a cell-specific RNAi (csRNAi) strategy for selective knockdown of *dyn-1* in DD neurons (Figure 4D). Removal of ventral SNB-1::GFP was impaired in *dyn-1(csRNAi)-*treated DD neurons thus suggesting that dynamin normally functions in DD neurons to promote presynaptic disassembly (Figure 4D).

Because CaN-dependent activation of Dynamin involves the dephosphorylation of residues in a proline-rich C-terminal “phospho-box” domain, we searched the phospho-proteome of *C. elegans* to identify DYN-1 amino acid residues that are phosphorylated *in vivo* (Zielinska et al., 2009). This approach detected three phosphorylated DYN-1 sites (S53, Y55, S59) in a proline-rich C-terminal domain (Figure 4E). To determine if the dephosphorylation state of DYN-1 modulates its role in DD remodeling, we forced expression of a phospho-resistant mutant (S53A, Y55A and S59A) (DYN-1AAA::SL2::2xNLS::TagRFP) in DD neurons, which our model predicts should accelerate the rate of presynaptic disassembly (Figure 4A). This experiment showed that ectopic expression of the DYN-1AAA results in precocious removal of presynaptic SNB-1::GFP from remodeling DD neurons (Figure 4F,G). In contrast, over-expression of wild-type DYN-1 (DYN-1::SL2::2xNLS::TagRFP) or DYN-1(OE) does not enhance the elimination of SNB-1::GFP from DD synapses (Figure 4G). These results confirm that dynamin can act cell-autonomously in DD neurons and that its dephosphorylation accelerates removal of DD synapses. Thus, our findings are consistent with the idea that CaN dephosphorylates dynamin and additional dephosphins to drive presynaptic disassembly in remodeling DD neurons. Because ADBE depends on CaN-dependent activation of dynamin, our results also substantiate the hypothesis that a bulk endocytic mechanism drives DD synaptic remodeling. Importantly, genes that are involved in other clathrin-independent endocytic mechanisms, (e.g., caveolin, Arf6) are dispensable for synapse elimination of GABA neurons (Mayor et al., 2014) (Figure S4D,E). Together, these results suggest that an ADBE-like mechanism drives DD remodeling. In contrast to the canonical function of ADBE of local recycling to sustain neurotransmission, however, our results suggest that a bulk endocytic mechanism effectively dismantles the DD presynaptic domain.

### Syndapin functions in a common pathway with UNC-8 to remove synaptic vesicle components in remodeling DD neurons

The dephosphin, Syndapin interacts with the plasma membrane via its F-BAR domain and also recruits downstream effectors through its SH3 domain (Figure 5A). During ADBE, Syndapin and Dynamin form a complex at the plasma membrane to promote the polymerization of branched-actin networks for endocytosis (Anggono et al., 2006; E. L. Clayton et al., 2009; Kessels & Qualmann, 2004). Consistent with the idea that Syndapin also functions with Dynamin to drive synaptic remodeling, we observed that RNAi knockdown of *sdpn-1/syndapin* in *unc-55* mutants impairs the removal of SNB-1::GFP from ventral GABAergic synaptic terminals (Figure S4D). Similarly, the elimination of endogenous GFP::RAB-3 from ventral neurites of DD neurons is disrupted in *sdpn-1* mutants (Figure 5B). Notably, removal of GFP::RAB-3 from remodeling DD synapses is impaired in *unc-8* mutants (Figure 5B)(Miller-Fleming et al., 2016, 2021) as predicted by our hypothesis that *unc-8* triggers a downstream bulk endocytic mechanism to dismantle the presynaptic domain (Figure 1A and Figure 4A). To test the idea that SDPN-1 functions in a common pathway with UNC-8, we created an *unc-8; sdpn-1* double mutant. This experiment confirmed that the ectopic retention of GFP::RAB-3 puncta in DD neurons of *unc-8; sdpn-1* double mutants is not significantly different from that of either the *unc-8* or *sdpn-1* single mutants (Figure 5B). Together, our findings argue that UNC-8 and SDPN-1 act in a common genetic pathway to remove synaptic vesicles in remodeling DD neurons.

**Figure 5.**
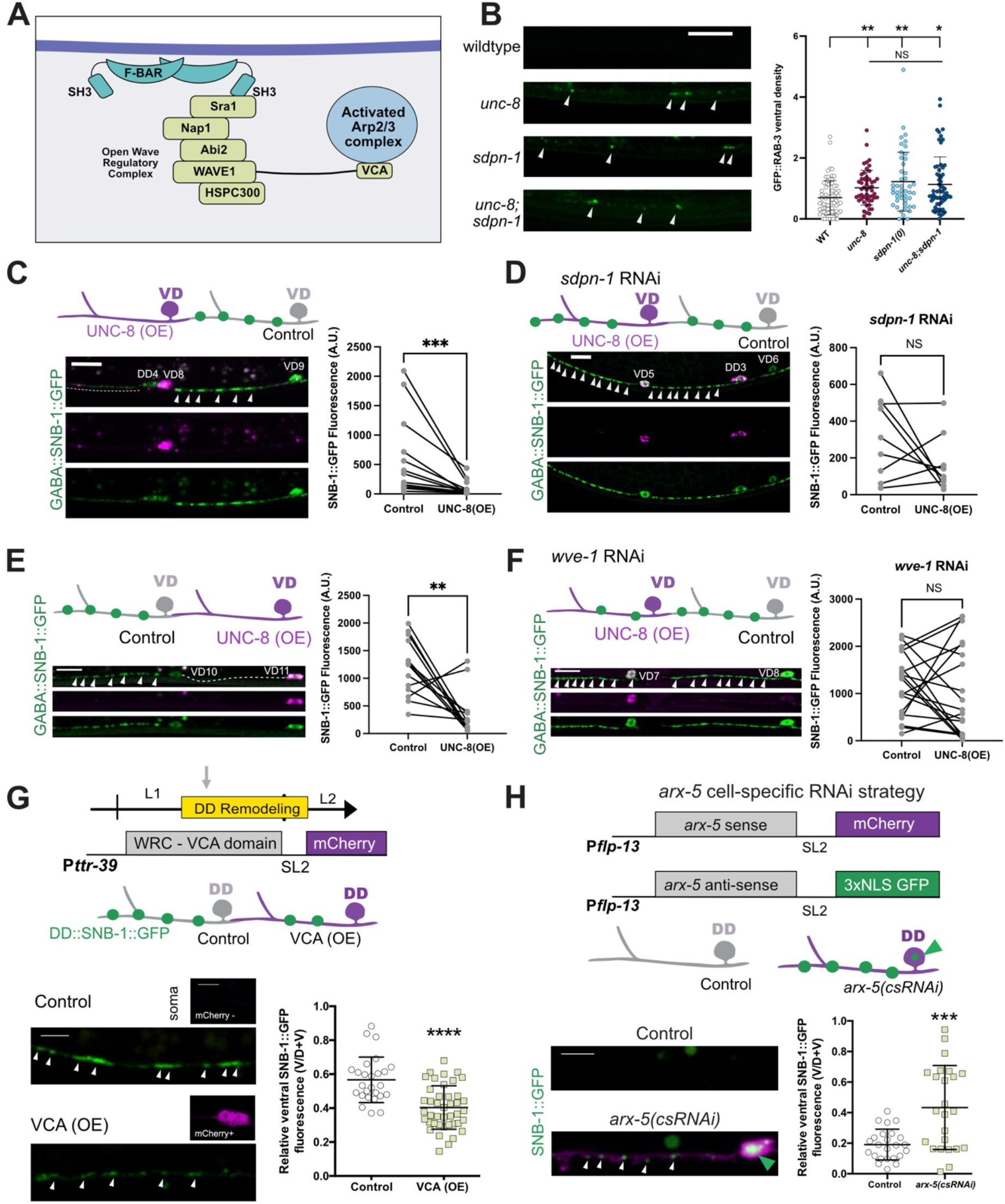
Branched-actin polymerization functions downstream of UNC-8 to remodel DD boutons. **A**. F-BAR proteins associated with the plasma membrane, like Syndapin (cyan), recruit the Wave Regulatory Complex (WRC) via SH3 domains to activate Arp2/3 for branched-actin polymerization. WRC components Sra1/GEX-2, Nap1/GEX-3, Abi2/ABI-1, WAVE1/WVE-1 and HSPC300/Y57G11C.1147 are conserved in *C. elegans*. The VCA domain (V: Verprolin homology domain, C: Central domain and A: Acidic domain) of WAVE1 interacts with Arp2/3 to promote its activation. **B**. SDPN-1 and UNC-8 function in a common genetic pathway. (Left) Residual endogenous GFP::RAB-3 puncta (arrowheads) in ventral DD neurites in wild type, in *unc-8* and *sdpn-1* single mutants and in *unc-8; sdpn-1* double mutants in early L4 larvae. (Right) Ventral GFP::RAB-3 density (puncta/10 micron) is not significantly different among *unc-8* (1.02 ± 0.5, n=51), *sdpn-1* (1.22 ± 0.9, n=46) and *unc-8; sdpn-1* double mutants (1.13 ± 0.9, n=62) mutants but is elevated in comparison to wild type (0.69 ± 0.6, n=73). **C**. (Left) Control VD neurons (gray) with anteriorly placed SNB-1::GFP puncta (arrowheads) vs UNC-8(OE) VD cells (magenta) in which SNB-1::GFP puncta are reduced (dashed line). (Right) Paired analysis of neighboring cells [control vs unc-8(OE)]: Control VDs (608 ± 688 A.U., n=13) and UNC-8(OE) VD neurons (94.9 ± 129 A.U., n=13). Wilcoxon matched-pairs signed rank test, *** p = 0.0002. L4 larvae were grown on empty vector bacteria as feeding RNAi control for Panel D. **D**. (Left) Treatment with *sdpn-1* RNAi results in retention of SNB-1::GFP puncta (arrowheads) in UNC-8(OE) VD neurons. (Right) Paired analysis of neighboring cells [control vs *unc-8(OE*)] in *sdpn-1* RNAi animals: Control VDs (321 ± 266 A.U., n=9) and UNC-8 (OE) VD neurons (163 ± 156 A.U., n=9). Wilcoxon matched-pairs signed rank test, p = 0.25. NS is Not Significant. Scale bar = 10 μm. L4 larvae. **E**. (Left) Control VD neurons (gray) with anteriorly placed SNB-1::GFP puncta (arrowheads) and UNC-8(OE) VD cells (magenta) with fewer SNB-1::GFP puncta (dashed line). (Right) Paired analysis of SNB-1::GFP fluorescence in neighboring cells [Control vs UNC-8(OE)]: Control VDs (1197 ± 507 A.U., n=14) and UNC-8 (OE) VD neurons (342 ± 398 A.U., n=14). Scale bar = 10 μm. L4 larvae. Animals grown on empty vector bacteria as feeding RNAi control for Panel D. **F**. (Left) Control VD neurons (gray) with anteriorly placed SNB-1::GFP puncta (arrowheads) and UNC-8(OE) VD cells (magenta) that maintain SNB-1::GFP puncta in (arrowheads) animals treated with *wve-1* RNAi. (Right) Paired analysis of SNB-1::GFP fluorescence in neighboring cells [Control vs UNC-8(OE)] in *wve-1* RNAi*-*treated animals: Control VDs (1194 ± 669 A.U., n=19) and UNC-8 (OE) VD neurons (1048 ± 969 A.U., n=19). Scale bar = 10 μm. L4 larvae. **G**. (Top) *Pttr-39* drives over-expression of the WRC-VCA domain [VCA(OE)] and cytosolic mCherry (magenta) in DD neurons. Remodeling is tracked with DD::SNB-1::GFP (green) during remodeling in the L1 stage (arrow). (Bottom Left) Representative images of ventral SNB-1::GFP puncta (arrowheads)in VCA(OE) DD neurons and control DD neurons. Scale bar = 5 μm. (Bottom RIght) Proportion of overall SNB-1::GFP fluorescence on the ventral side of control DDs (0.56 ± 0.1, n=26) and VCA(OE) (0.40 ± 0.1, n=42) cells during remodeling at the late L1 stage. Unpaired t-test, **** < 0.0001. **H**. (Top) Cell specific RNAi (csRNAi) of *arx-5* in DD neurons with *arx-5* sense and antisense transcripts each co-expressed with either cytosolic mCherry or nuclear-localized GFP. Non-fluorescent DD neurons serve as controls (gray) whereas DDs with nuclear GFP (green arrowhead) and cytosolic mCherry correspond to *arx-5(csRNAi)* knockdown cells. (Bottom, Left) Representative images of ventral SNB-1::GFP puncta (white arrowheads) in control and *arx-5(csRNAi)* DD cells in L4 larvae. (Bottom, Right) Proportion of overall SNB-1::GFP fluorescence on the ventral side in control DDs (0.19 ± 0.1, n=24) and *arx-5(csRNAi)* (0.43 ± 0.3, n=24) cells of L4 larvae. Unpaired t-test, **** < 0.0001. Scale bar = 10 μm.

Our model also predicts that SDPN-1 functions downstream of UNC-8 to drive presynaptic disassembly (Figure 4A). To test this idea, we asked if SDPN-1 is required for the ectopic removal of ventral GABAergic synapses arising from forced expression of UNC-8 in VD neurons. As noted above, in the wild type, VD neurons do not express UNC-8 and normally establish stable GABAergic synapses with ventral muscles (Miller-Fleming et al., 2016; White et al., 1986). UNC-8 over-expression or UNC-8(OE) in VD neurons, however, is sufficient to remove SNB-1::GFP from ventral VD presynaptic domains (Figure 4C) (Miller-Fleming et al., 2016, 2021). This effect is abrogated, however, by RNAi knockdown of *sdpn-1* (Figure 5D) thus supporting the hypothesis that SDPN-1 acts downstream of UNC-8 for synapse elimination. Consistent with a presynaptic role for SDPN-1 for DD synaptic disassembly, SDPN-1::mCherry is closely associated with the presynaptic marker SNB-1::GFP during the remodeling window (Figure S5A-B). To summarize, results in this section are consistent with the proposal that UNC-8 drives synaptic disassembly in a pathway that involves the F-BAR protein SDPN-1/Syndapin and thus support the idea that a bulk endocytic mechanism resembling ADBE (Figure 4A and 5A) promotes synapse elimination in remodeling DD neurons.

### Branched actin polymerization drives DD synaptic remodeling

Close association of F-BAR domains with the plasma membrane mobilizes branched-actin activators to initiate membrane invagination during endocytosis (Kessels & Qualmann, 2004). Because the F-BAR protein SDPN-1/Syndapin promotes branched actin polymerization via its SH3 domain (Kessels & Qualmann, 2004), we conducted a candidate screen of known actin-related downstream effectors for roles in synaptic remodeling. For this purpose, we utilized the *unc-55* paradigm (Figure S5C-D) to determine that RNAi knockdown of components of the Arp2/3 complex and the Wave Regulatory Complex (WRC) impairs the removal of SNB-1::GFP from ventral synapses in remodeling GABAergic neurons (Figure S5F) (Petersen et al., 2011). These findings are consistent with the hypothesis that SDPN-1/Syndapin drives synapse elimination by recruiting downstream effectors of actin polymerization, the WRC and Arp2/3 complex, to remodeling GABAergic synapses (Figure 5A). Intriguingly, an additional Arp2/3 activator, N-WASP, is dispensable for this process (Figure S5F).

Additional evidence suggests that the WRC acts downstream of UNC-8. RNAi knockdown of the WRC component, *wve-1*, impairs the removal of SNB-1::GFP from the ventral synapses of VD neurons that over-express UNC-8 (Figure 5E-F). Thus, our results are consistent with the idea that UNC-8 drives synapse removal by activating branched actin polymerization via the F-BAR protein, SDPN-1/Syndapin and the WRC. For endocytosis, F-BAR proteins interact with the WRC which in turn deploys its VCA domain to activate the Arp2/3 complex (Chen et al., 2010; Girao et al., 2008; Ismail et al., 2009; Mooren et al., 2012) (Figure 5A). Because the VCA domain of WRC is sufficient to activate the Arp2/3 complex *in vitro* (Machesky et al., 1999), we over-expressed the VCA domain in DD neurons to test the prediction that elevated VCA activity would accelerate DD remodeling. Remodeling DD neurons with forced VCA expression or VCA(OE) showed fewer ventral SNB-1::GFP puncta than control DD neurons that did not express the VCA domain (Figure 5G). These results support the hypothesis that the WRC activates branched-actin polymerization in remodeling DD neurons to promote disassembly of the presynaptic apparatus.

To track the Arp2/3 complex during remodeling, we GFP-labeled the conserved Arp2/3 component, ARX-5/p21, for dual-color imaging with the synaptic vesicle marker mCherry::RAB-3. Airyscan imaging during the remodeling window detected ARX-5::GFP puncta closely associated with RAB-3::mCherry at remodeling ventral DD presynaptic boutons (Figure S5G). To rigorously test the hypothesis that branched-actin polymerization is required in DD neurons to promote presynaptic disassembly, we used cell specific RNAi (csRNAi) to achieve DD-specific knockdown of *arx-5* (Figure 5H). Importantly, ARX-5/p21 is required for Arp2/3 function as shown by the finding that selective depletion of the ARX-5 subunit decreases the rate of Arp2/3-dependent actin polymerization (Gournier et al., 2001). We observed significant retention of SNB-1::GFP puncta at the presynaptic terminals of *arx-5(csRNAi)*-treated DD neurons in comparison to control cells, indicating the failure of SNB-1::GFP elimination in *arx-5*-depleted cells (Figure 5H). Thus, our results point to a cell autonomous role for Arp2/3 and branched actin polymerization in the removal of presynaptic components of remodeling DD neurons.

### Actin polymerization and transient RAB-3 particles are elevated during DD synaptic remodeling and depend on UNC-8

Our genetic studies show that components that drive branched-actin polymerization promote removal of DD synapses (Figure 5). These findings predict that actin polymerization is upregulated during DD remodeling as presynaptic components are dismantled. For a direct test of this idea, we labeled the actin cytoskeleton with LifeAct::mCherry (Riedl et al., 2008) and synaptic vesicles with endogenous GFP::RAB-3 in DD neurons. Dual-color live cell imaging detected GFP::RAB-3 puncta surrounded by LifeAct::mCherry-labeled actin (Figure 6A-B). Actin dynamics at synapses were recorded by time-lapse imaging as changes in LifeAct::mCherry fluorescence in regions that corresponded to GFP::RAB-3-labeled clusters (Figure 6C-G) (See Methods). Kymograph analysis revealed robust elevation of both LifeAct::mCherry fluorescence intensity and dynamics at presynaptic regions during the remodeling window (Figure 6C-E and Figure S6A-D). To quantify this effect, we calculated the standard deviation of all synaptic LifeAct::mCherry traces and determined that fluctuations in LifeAct::mCherry fluorescence are enhanced during the remodeling window in comparison to an earlier developmental period before remodeling (Figure S6B-D) (Video S4 and S5). Interestingly, live imaging detected instances of high actin dynamics in axonal regions with mobile GFP::RAB-3 particles (Figure 6C, arrows) (Video S5). These dynamic GFP::RAB-3 puncta are smaller than nearby, more stable GFP::RAB-3 clusters from which they appear to arise based on their diagonal trajectories (Figure 6C, arrows). Dynamic and small GFP::RAB-3 puncta also appear short-lived as they can spontaneously disappear from the intersynaptic space (Figure 6C, arrowhead). Quantification revealed that transient GFP::RAB-3 particles predominate during the remodeling window but are detected less often before DD remodeling ensues (Figure S6E). Importantly, we observed instances in which the disappearance of a transient GFP::RAB-3 particle correlates with temporal elevation of LifeAct::mCherry (Figure 6C’’) (Video S6). Notably, increased actin dynamics (Figure S6D) and transient GFP::RAB-3 puncta (Figure S6E) are correlated with an overall reduction in the density of RAB-3::GFP puncta (Figure S6F). Together, these results suggest that both actin dynamics and transient GFP::RAB-3 puncta are upregulated during DD remodeling when presynaptic domains are disassembled.

**Figure 6.**
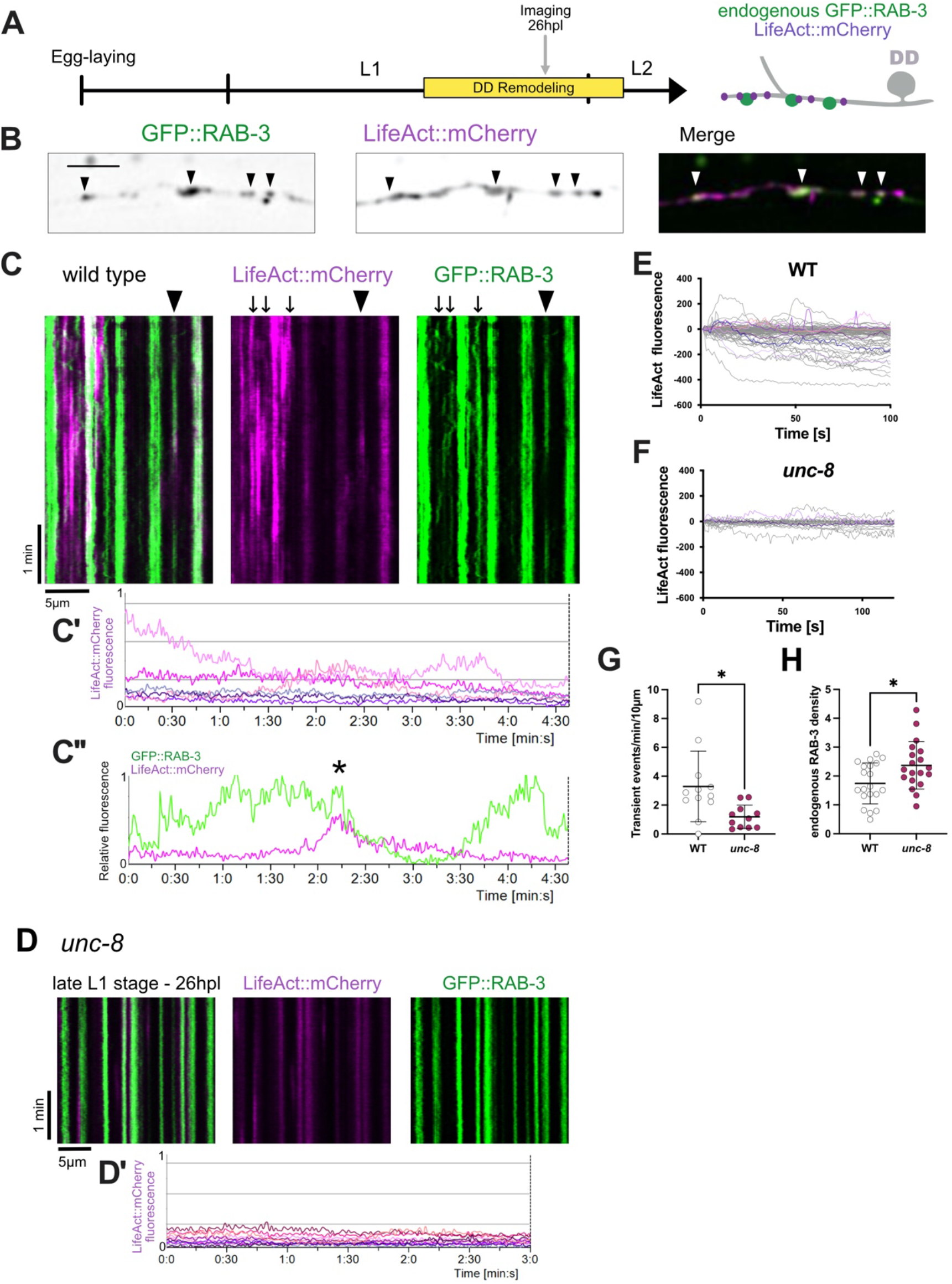
Actin polymerization is upregulated during DD remodeling and correlates with RAB-3 dynamics. **A**. Ventral regions of DD neurons were imaged during remodeling (gray arrow) to monitor actin dynamics with LifeAct::mCherry (magenta) and synaptic vesicles with endogenous GFP::RAB-3 (green) in wild-type and *unc-8* mutant animals. **B**. Live-imaging reveals endogenous GFP::RAB-3 (green) associated with LifeAct::mCherry (magenta). Arrowheads point to locations of GFP::RAB-3 clusters. Scale Bar = 5 μm. **C-F**. Elevated actin dynamics during DD remodeling in wild-type animals but not in *unc-8* mutants. **(C)** During remodeling, LifeAct::mCherry associates with both stable and transient GFP::RAB-3 puncta in wild-type animals. Arrows and arrowhead (see below) point to locations of dynamic GFP::RAB-3 puncta. **(C’)** Line scans from kymograph show dynamic LifeAct::mCherry fluorescence during remodeling, n = 6 boutons from kymograph depicted in C. **(C’’)** Line scans of LifeAct::mCherry and GFP::RAB-3 from kymograph trace (arrowhead) showing reduction in GFP::RAB-3 signal with transient elevation of LifeAct::mCherry fluorescence (asterisk). **(D)** Kymographs depict actin polymerization (LifeAct::mCherry) and stable GFP::RAB-3 puncta in *unc-8* mutants. **D’** Line scans from kymograph showing LifeAct::mCherry fluorescence at GFP::RAB-3 boutons (N = 11 boutons from kymograph depicted in D). The Y-axis is normalized to the maximum value of C’ line scans. **(E-F)** LifeAct::mCherry fluorescence from GFP::RAB-3 regions of all videos analyzed (normalized to t_0_ = 0) in **(E)** wild type (n = 12 videos) and **(F)** *unc-8* mutants (n = 11 videos). **G**. Fewer transient GFP::RAB-3 events were detected in *unc-8* mutants (1.2 ± 0.8, n = 11 videos) compared to WT (3.2 ± 2.4, n = 12 videos). T-test, * is p<0.05. **H**. *unc-8* mutants retain more GFP::RAB-3 puncta (2.37± 0.8, n = 19 snapshots) than wild type animals (1.74 ± 0.7, n = 20 snapshots) during the remodeling window (L2 larval stage). T-test, * is p<0.05.

Because our results suggest that UNC-8 functions upstream of the WRC (Figure 5E, F) which promotes branched actin polymerization through the activation of the Arp2/3 complex (Figure 5A), we next asked if actin polymerization is defective in *unc-8* mutants. Our *in vivo* analysis detected a striking reduction in LifeAct::mCherry dynamics during the DD remodeling period in *unc-8* mutant L1 larvae in comparison to wild-type controls (Figure 6D-F) (Video S7). Mobile particles labeled with endogenous GFP::RAB-3 were also reduced in remodeling DD neurons relative to wild type in *unc-8* mutants (Figure 6G). Notably, these defects are correlated with increased density of residual ventral GFP::RAB-3 puncta in *unc-8* mutants during remodeling (Figure 6H). Together, these findings are consistent with the hypothesis that UNC-8 normally promotes actin polymerization to accelerate removal of GFP::RAB-3 from remodeling DD synapses.

### DEG/ENaC-dependent endocytic vesicles populate remodeling DD neurons

During ADBE, actin polymerization at the plasma membrane aids the formation of bulk endosomes (Emma L Clayton & Cousin, 2009). We have shown that UNC-8/DEG/ENaC elevates Ca^2+^ at the synapse (Figure 2), and that key elements of the ADBE mechanism including branched actin polymerization function downstream of UNC-8 (Figure 5 and 6) (Miller-Fleming et al., 2016). These findings predict that UNC-8 activity promotes the formation of bulk endosomes in remodeling DD neurons. To test this hypothesis, we used serial section electron microscopy (EM) to generate 3D-reconstructions of the ventral neurites of remodeling DD neurons at the late L1 larval stage in the wild type and in an *unc-8* mutant (Figure 7A) (Video S8). We annotated key presynaptic structures including synaptic vesicles, dense core vesicles, dense projections (e.g., presynaptic densities) in both wild type and *unc-8* (Figure S7A-D). We also noted large, clear, circular endocytic structures that resemble bulk endosomes (Figures 7A-E and S7C) (Emma L Clayton & Cousin, 2009). These endocytic structures were visible in the DD neurites of > 30% of EM sections or profiles in the wild type but in only ∼15% of profiles in the *unc-8* mutant and this difference is statistically significant (Figure 7E). Interestingly, the average volume of endocytic structures in *unc-8* (41.2 ± 5.2 × 10^4^ nm^3^) was significantly larger than in the wild type (22.5 ± 3.4 × 10^4^ nm^3^) (Figure S7C). Our determination that ventral DD dendrites in *unc-8* mutants contain fewer large endocytic structures than in the wild type offers strong support for the hypothesis that UNC-8 promotes the formation of bulk endosomes in remodeling DD neurons.

**Figure 7.**
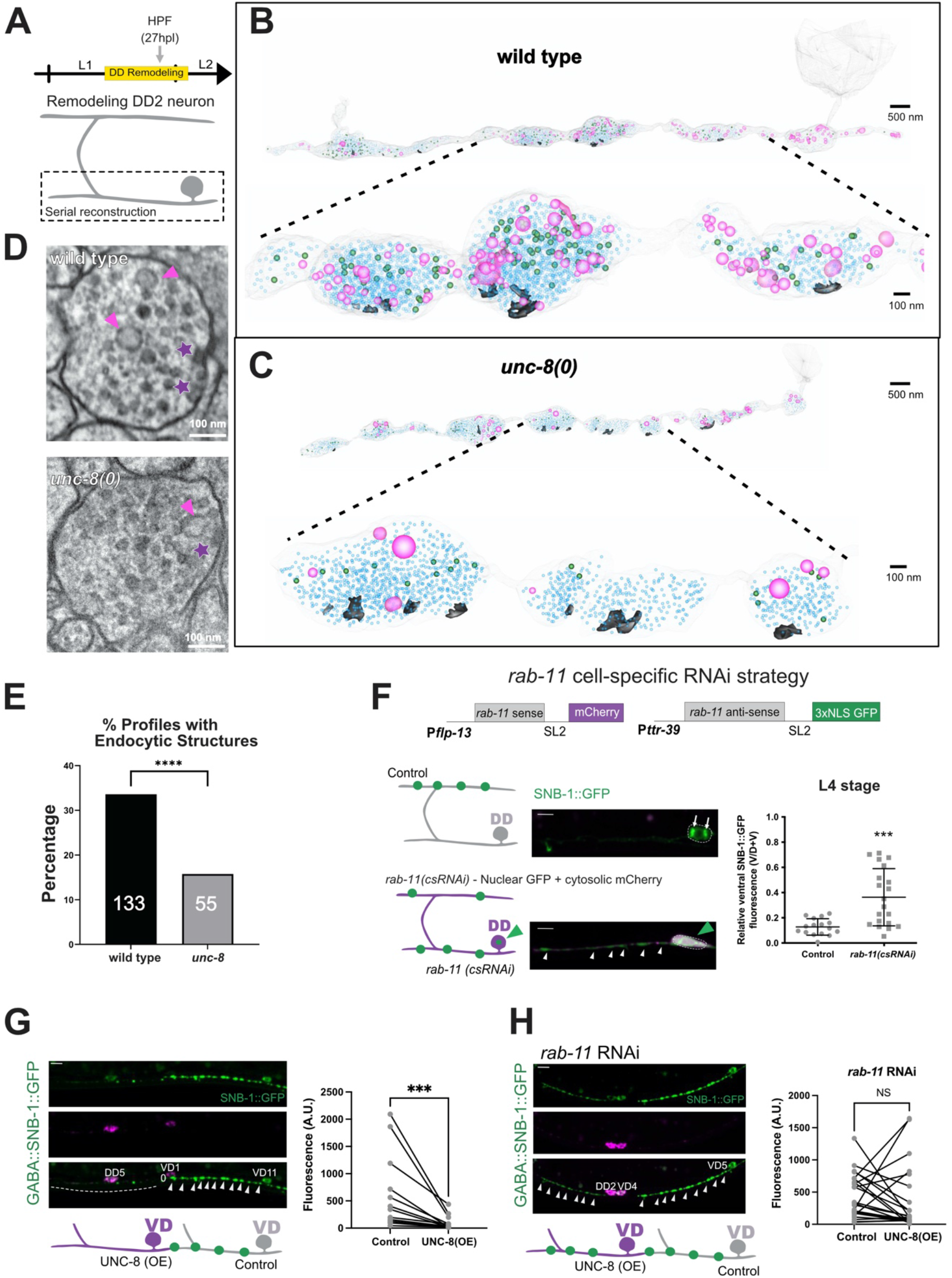
The recycling endosome component, RAB-11, is required for UNC-8-dependent synapse elimination. **A**. L1 stage larvae were subjected to High Pressure Freezing (HPF) during the DD remodeling period (27hpl) and serial sections were imaged in the electron microscope. **B-C**. Full reconstructions of **(B**) wild type and (**C**) *unc-8(0)* DD2 neurons with enlargements below. Neuronal membrane (transparent white), Synaptic Vesicles (SVs) (light blue spheres), Dense Core Vesicles (DCVs) (green spheres), dense projections (grey), and endocytic structures (magenta). **D**. Representative TEM sections containing presynaptic densities (asterisk) and endocytic structures (arrowhead) from reconstructed wild-type (top) and *unc-8(0)* (bottom) DD2 neurons. **E**. Percentage of TEM profiles (sections) containing endocytic structures in reconstructed DD2 neurons in wild type and *unc-8(0)*. Numbers (white) denote sections with endocytic structures for each genotypes. Statistical significance was calculated with Fischer’s Exact Test (**** is p<0.00001). **F**. Cell-specific RNAi of *rab-11* retards removal of GFP::RAB-3 from remodeling DD neurons. (Top) *rab-11* cell-specific RNAi strategy. (Middle) Schematic of L4 stage control DD neuron (gray) with SNB-1::GFP puncta in the dorsal cord vs (Bottom) *rab-11(scRNAi)-*treated DD neuron (magenta) with *rab-11* sense and antisense transcripts each co-expressed with either cytosolic mCherry or nuclear-localized GFP (green arrowhead) and residual SNB-1::GFP puncta in the ventral cord. Representative images at the L4 larval stage show (Middle) depletion of ventral SNB-1::GFP puncta in the wild type (arrows denote residual SNB-1::GFP in Golgi) vs (Bottom) retention of ventral SNB-1::GFP in *rab-11(csRNAi)*-treated DD neurons. Dashed lines demarcate DD cell soma. Scale bar = 10 μm. (Right) Proportion of overall SNB-1::GFP fluorescence on the ventral side in control (0.13 ± 0.1, n = 16) vs *rab-11(csRNAi)*-treated (0.36 ± 0.2, n = 20) DD neurons. Mann-Whitney test, *** p = 0.0003. **G**. (Left) Control VD neurons (VD11, gray) with anteriorly placed SNB-1::GFP puncta (arrowheads) and UNC-8(OE) VD cells (VD10, magenta) with fewer SNB-1::GFP puncta in anterior region (dashed line). (Right) Paired analysis of ventral SNB-1::GFP fluorescence in neighboring cells, Control VDs (607 ± 688 A.U., n=13) vs UNC-8 (OE) VD neurons (94.8 ± 129 A.U., n=13). Wilcoxon matched-pairs signed rank test, *** p = 0.0002. Scale bar = 10 μm. **H**. (Left) Anteriorly placed SNB-1::GFP puncta (arrowheads) in Control VD neurons (VD5, gray) and UNC-8 (OE) VD cells (magenta) treated with *rab-11* RNAi. Scale bar = 10 μm. (Right) Paired analysis of ventral SNB-1::GFP fluorescence in neighboring cells, Control VDs (439 ± 341 A.U., n=23) and UNC-8(OE) VD neurons (360 ± 500 A.U., n=23). Wilcoxon matched-pairs signed rank test, p = 0.211. NS is Not Significant.

### Recycling endosomes function downstream of UNC-8 to mediate synapse elimination and dorsal assembly

As noted above, the canonical ADBE mechanism generates large bulk endosomes that capture presynaptic membrane and SV components for recycling at the adjacent synapse (Cheung & Cousin, 2019). Our finding of abundant endocytic structures in remodeling DD axons and their dependence on UNC-8 which promotes presynaptic disassembly, suggested that SV components could be recycled by an ADBE-like mechanism for reassembly, in this case, to nascent presynaptic domains in dorsal DD neurites (Figure 1A)(Park et al., 2011).

A mechanism for distal recycling of presynaptic components would likely require a transport system involving intracellular vesicles or endosomes (Ivanova & Cousin, 2022). Rab11 is a small GTPase required for recycling endosome function (Ullrich et al., 1996). We adopted two approaches to test the idea that RAB-11 promotes remodeling of DD presynaptic terminals. First, we used cell-specific RNAi to knockdown *rab-11* in remodeling DD neurons. We observed that ventral SNB-1::GFP fluorescence in L4 larvae was elevated in *rab-11(csRNAi)-*treated DD neurons in concert with a reciprocal decrease in dorsal SNB-1::GFP signal. In contrast, control DD cells did not retain ventral SNB-1::GFP and successfully assembled SNB-1::GFP puncta in the dorsal cord by the L4 stage (Figure 7F). These findings support the notion that *rab-11* and thus recycling endosomes are required for the removal of synaptic proteins from ventral terminals as well as for their assembly at dorsal synapses. To determine if RAB-11 functions downstream of UNC-8, we asked if UNC-8-dependent synapse removal in VD neurons (Figure 7G-H) requires RAB-11. In this experiment, global RNAi knockdown of *rab-11* was sufficient to prevent UNC-8(OE)-dependent elimination of transport disassembled ventral synapses to sites of nascent presynaptic assembly in dorsal DD neurites, a result consistent with the hypothesis that RAB-11 normally functions downstream of UNC-8/DEG/ENaC to promote DD presynaptic remodeling.

In a second experiment to test the idea that recycling endosomes are involved in DD synaptic remodeling, we designed a two-color labeling strategy to track the recycling endosome marker RAB-11::TagRFP and the synaptic vesicle-associated component, GFP::RAB-3, in remodeling DD neurons (Figure S7E). Live-imaging detected dynamic RAB-11::TagRFP particles that actively traffic along both ventral and dorsal DD neurites during the remodeling window (Figure S7F-G). RAB-11::TagRFP particles were also observed entering the commissure that connects the ventral and dorsal DD neurites (Video S9). Subsets of RAB-11::TagRFP particles are closely apposed to stable GFP::RAB-3 puncta (Figure S7F-G) with striking examples of dynamic association and dissociation with the periphery of GFP::RAB-3 puncta (Figure S7F-G, yellow arrowheads). Intriguingly, in some cases, this association of dynamic RAB-11::TagRFP particles is correlated with the disappearance of otherwise stable GFP::RAB-3 puncta (Figure S7G).

### RAB-11 promotes recycling of ventral RAB-3 to nascent, dorsal synapses in remodeling DD neurons

Having shown that *rab-11(csRNAi)* impairs DD remodeling and that RAB-11::TagRFP-labeled particles are actively trafficked in both dorsal and ventral DD neurites, we next performed an experiment to ask if recycling of ventral components to nascent dorsal synapses depends on RAB-11. First, we designed a flippase-dependent cassette at the N-terminus of the *rab-3* locus that encodes the photoconvertible fluorescent protein Dendra-2 (Figure 8A) and used a DD-specific promoter (*Pflp-13*) to drive expression of the resultant endogenous Dendra-2::RAB-3 in DD neurons. Exposure to UV light induces an irreversible green-to-red change in the Dendra-2 fluorescence emission (Figure 8B). We used this approach to photoconvert Dendra-2::RAB-3 at ventral DD synapses (green) with UV light before the onset of remodeling in the L1 stage (18-20 hpl) and to track photoconverted Dendra-2::RAB-3 (red) during the remodeling period in the late L1 (28 hpl) and after remodeling is complete in the early L3 larval stage (38 hpl) (Figure S8A-D). Imaging during the remodeling window detected photoconverted Dendra-2::RAB-3 in both ventral and dorsal nerve cords. Later, by the L3 stage, photoconverted Dendra-2::RAB-3 had been efficiently removed from the ventral side with > 60% recovered at dorsal DD neurites. (Figure S8E). These findings confirm an earlier report that Dendra::RAB-3 over-expressed from a transgenic array is also recycled from ventral to dorsal synapses in remodeling DD neurons (Park et al., 2011).

**Figure 8.**
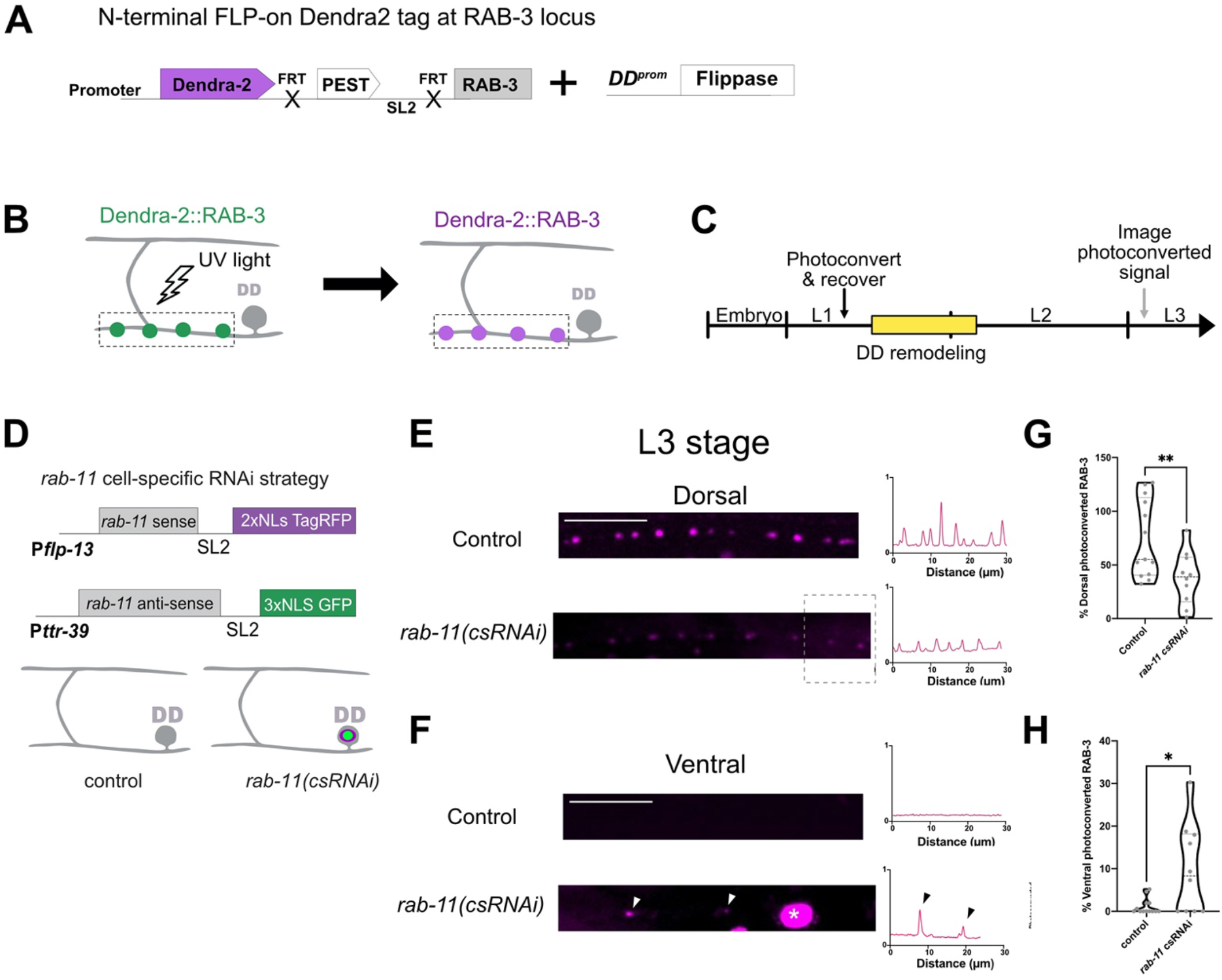
RAB-11 promotes recycling of RAB-3 from old to new presynaptic boutons in remodeling DD neurons. **A**. Endogenous labeling of RAB-3 with photoconvertible protein Dendra-2 in DD neurons. *pflp-13* drives flippase in DD neurons to fuse Dendra-2 to the N-terminus of the endogenous RAB-3 protein.**B**. Schematic of Dendra-2::RAB-3 photoconversion. UV irradiation of ventral Dendra-2::RAB-3 in individual DD neurons (dashed box) produces photoconverted Dendra-2::RAB-3 puncta (magenta). **C**. Experimental set up: Dendra-2::RAB-3 was photoconverted before DD remodeling and imaged after remodeling at the L3 larval stage (gray arrow). **D**. Cell specific RNAi (csRNAi) knockdown of rab-11 in DD neurons is achieved by co-expression of *rab-11* sense transcripts with nuclear-localized marker, 2xNLS TagRFP and *rab-11* anti-sense transcript with nuclear-localized marker, 3xNLS GFP. Non-fluorescent DD neurons (gray) serve as controls whereas DD neurons with nuclear-localized TagRFP and GFP (magenta + green) correspond to *rab-11(csRNAi)* knockdown cells. **E-F. (**Left) Representative images (L3 larval stage) of photoconverted Dendra-2::RAB-3 (magenta) in control and *rab-11(csRNAi)* DD neurons in dorsal (top) and ventral (bottom) DD neurites. Arrowheads denote ventral retention of photoconverted Dendra-2::RAB-3 in *rab-11(csRNAi)* neurons and asterisk labels nuclear-localized TagRFP expressed in *rab-11(csRNAi)* DD cells. (Right) Line scans of photoconverted Dendra-2::RAB-3 dorsal (top) and ventral (bottom) DD neurites of control and *rab-11(csRNAi)* knockdown cells. Arrowheads denote ventral retention of photoconverted Dendra-2::RAB-3 in *rab-11(csRNAi)* neurons. Scale bar = 10 μm. **G-H**. Percent of photoconverted Dendra-2::RAB-3 signal that recycles to the dorsal cord **(G)** or remains on the ventral nerve cord **(H)** at the L3 stage. **(G)** *rab-11(csRNAi)* DD cells recycle less photoconverted signal to the dorsal nerve cord (37.7 ± 24.8%, n=10) than controls (74.0 ± 36.2 %, n=13). Unpaired t-test, ** p = 0.0064. **(H)** *rab-11(csRNAi)* DD cells retain more photoconverted signal on the ventral side (9.96 ± 10.5 %, n=10) than controls (1.09 ± 1.8 %, n=13). Mann Whitney test, * p = 0.0269.

To determine if RAB-11 is required for RAB-3 recycling, we used cell-specific RNAi (csRNAi) to knock down *rab-11* in DD neurons. In this experiment, the cell nuclei of *rab-11(csRNAi)*-treated DD neurons are marked with GFP and tagRFP whereas neighboring control DD neurons that display unlabeled DD nuclei should express native levels of *rab-11* (Figure 8D). Dendra-2::RAB-3 was photoconverted before remodeling (18 hpl) and imaged after remodeling, at the L3 stage (38 hpl) (Figure 8C). In control DD neurons, photoconverted Dendra2::RAB-3 protein was efficiently removed from the ventral side and ∼70% recovered at nascent dorsal DD synapses (Figure 8E-G) as previously observed for the wild-type (Figure S8D-E). In contrast, *rab-11(csRNAi)-*treated DD neurons retained ventral photoconverted Dendra-2::RAB-3 signal and showed lower levels (∼40%) of recycled RAB-3 at dorsal synapses (Figure 8E,F, H). Together, these findings establish that RAB-3 protein, initially associated with ventral DD synapses, is recycled to nascent dorsal synapses in remodeling DD neurons in a mechanism that depends on the canonical recycling endosome component, RAB-11.

### UNC-8 promotes RAB-3 recycling for assembly at nascent dorsal synapses in remodeling DD neurons

Having established that UNC-8 promotes an endocytic mechanism for removing ventral presynaptic components, we hypothesized that an *unc-8* mutants would impair recycling of RAB-3 to dorsal DD synapses. To test this idea, we tracked photoconverted Dendra-2::RAB-3 from ventral to dorsal synapses (Figure 8A-C). This experiment confirmed that *unc-8* mutants show substantially reduced levels of recycled (i.e., photoconverted) Dendra-2::RAB-3 at dorsal synapses (Figure 9C). In an independent experiment, we measured the accumulation of endogenous GFP::RAB-3 puncta at dorsal synapses and observed a significant but weaker effect in *unc-8* mutants (Figure 9F,G). This disparity could mean that dorsal DD boutons are assembled from a combination of RAB-3 recycled from ventral synapses plus nascent RAB-3 that originates from the DD cell soma. In this model, the *unc-8* mutant impairs RAB-3 recycling but does not retard the incorporation of nascent RAB-3 at dorsal synapses (Miller-Fleming et al., 2016). Interestingly, by the L3 stage, photoconverted Dendra-2::RAB-3 is efficiently removed from ventral DD synapses in *unc-8* mutants (Figure 9D-E). In contrast, at an earlier timepoint during the remodeling window (L2 stage) residual ventral GFP::RAB-3 particles are elevated in *unc-8* mutants in comparison to wild type (Figure 6H). These results suggest that RAB-3 removal is delayed but not completely blocked in *unc-8* mutants perhaps due to the activity of an additional parallel-acting pathway that we have previously reported (Miller-Fleming et al., 2021). Taken together, our results suggest that UNC-8 promotes an endocytic mechanism that dismantles the presynaptic apparatus in ventral DD neurites for recycling at nascent DD synapses in the dorsal nerve cord.

**Figure 9.**
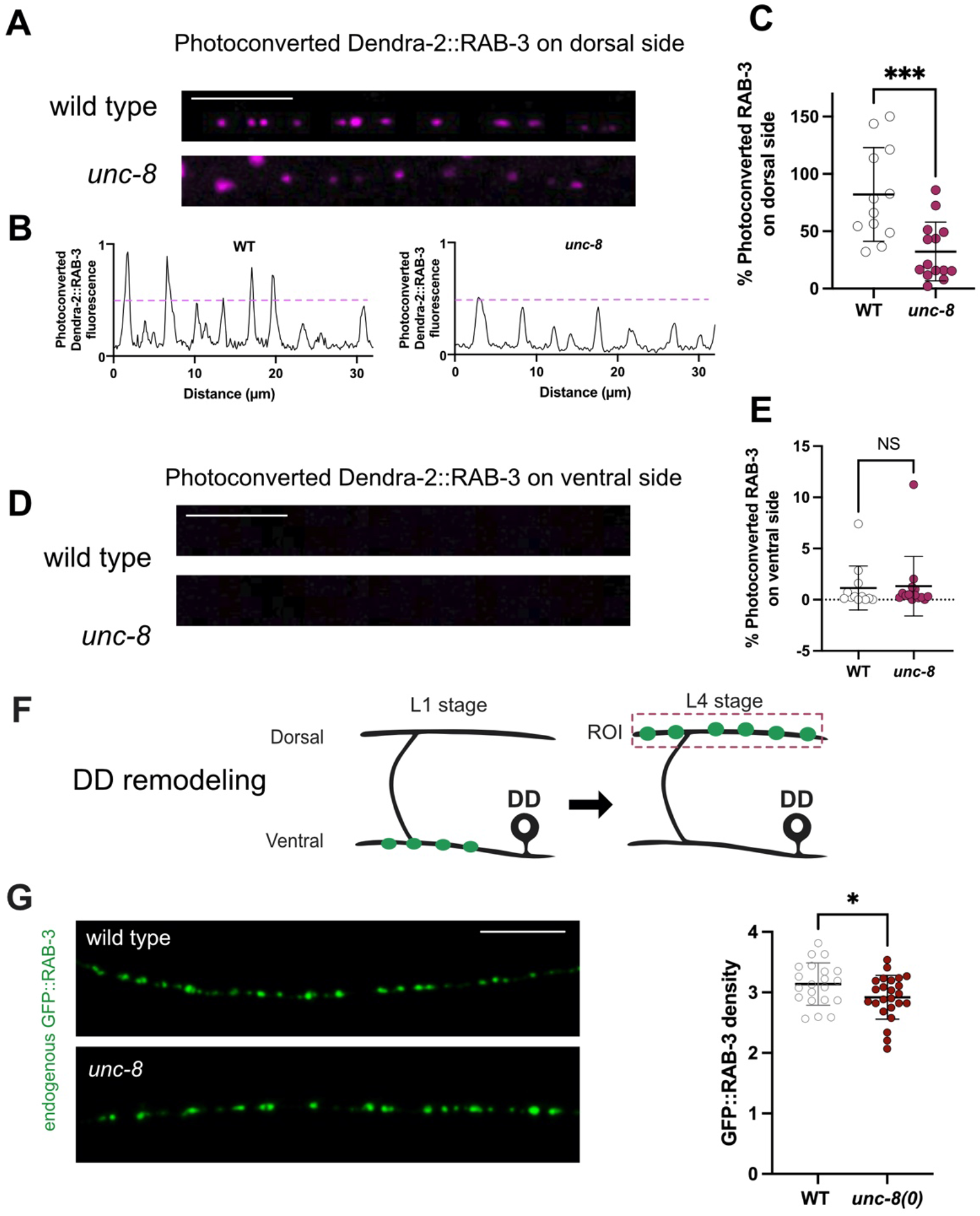
UNC-8 promotes recycling from ventral to dorsal DD boutons. **A**. Fluorescent images of photoconverted Dendra-2::RAB-3 in the dorsal nerve cord of (top) wild-type and (bottom) *unc-8* animals after the remodeling period in L3. Asterisk denotes autofluorescence. Scale bar = 10 μm. **B**. Normalized line scans across the dorsal cords of (left) wild-type (WT) and (right) *unc-8* mutant animals. All Y-axes are normalized to the brightest punctum in wild type. Dashed line shows threshold at 50% of the signal. **C**. *unc-8* (32.3 ± 25.6%, n=14) mutants recycle less photoconverted Dendra-2::RAB-3 to the dorsal nerve cord than wild type (WT) (82.0 ± 40.8%, n=12). Ordinary One-Way ANOVA with Dunnett’s multiple comparison test. *** p = 0.0009. **D**. Photoconverted Dendra-2::RAB-3 fluorescent signal is depleted in the ventral nerve cords of wild-type and *unc-8* animals at the L3 stage. Scale bar = 10 μm. **E**. Quantification confirms that a small fraction of photoconverted Dendra-2::RAB-3 signal retained in the ventral nerve cord at the L3 stage in wild type (WT) (1.14 ± 2.1%, n=12) and *unc-8* (1.32 ± 2.9%, n=14). Kruskal-Wallis with Dunn’s multiple comparison test against WT, NS is Not Significant *unc-8*, p = 0.96. **F**. Schematic of GFP::RAB-3 relocation from DD ventral synapses in L1 larvae to dorsal locations in the L4 stage (ROI = Region of Interest). **G**. UNC-8 promotes localization of GFP::RAB-3 at dorsal DD synapses. (Left) Representative images of GFP::RAB-3 puncta in dorsal DD neurites in the wild type and *unc-8* mutants at the L4 larval stage. (Right). Quantification of dorsal GFP::RAB-3 density (puncta/10 um) in the wild type (WT) (3.14 ± 0.36%, n=21) and *unc-8* mutants (2.91 ± 0.34%, n=24). unpaired T test, *p = 0.046.

## Discussion

Circuits are actively remodeled during development as new synapses are assembled while others are removed. The underlying mechanisms derive from the intersection of genetic and activity-dependent programs, but the cell biological drivers of circuit refinement are largely unknown. To address this question, we have exploited the ready accessibility of *C. elegans* for live cell imaging and genetic analysis to investigate developmentally-regulated synaptic remodeling in the DD class of GABAergic neurons (Cuentas-Condori & Miller, 2020; Kurup & Jin, 2016). We show that presynaptic domains are dismantled, and synaptic vesicle components recycled to new locations in a mechanism that involves both genetic and activity-dependent pathways. Our results demonstrate that the DEG/ENaC channel, UNC-8, is selectively expressed during a specific developmental period to promote presynaptic disassembly in a process involving the Ca^2+^-dependent phosphatase, Calcineurin/CaN, and its targets, the GTPase Dynamin and the F-BAR protein Syndapin (Figure 1B & 4A). Downstream effectors, the WAVE Regulatory Complex (WRC) and Arp2/3 (Figures 5E-H & S5F) promote actin polymerization (Figure S6D) to drive the relocation of the synaptic vesicle-associated protein, RAB-3, to new DD synapses (Figure 8). Underscoring the role of endocytic recycling in synaptic remodeling, we determined that RAB-11, a key component of recycling endosomes, is required for the translocation of RAB-3 to new presynaptic domains in a pathway that functions downstream of UNC-8 (Figure 8 & 7G). Moreover, we used EM reconstruction to show that UNC-8 promotes the appearance of abundant, large endocytic structures in remodeling DD neurites (Figure 8A-E). The mechanism that we have described is strikingly similar to that of Activity-Dependent-Bulk Endocytosis (ADBE) which functions during periods of high activity to recycle synaptic vesicles for reuse at adjacent synapses (Cousin, 2009). Our results suggest that the canonical ADBE mechanism can be repurposed during development for the radically different outcome of dismantling the presynaptic apparatus for reassembly at new locations.

### The DEG/ENaC channel UNC-8 drives a Ca^2+^-dependent mechanism of presynaptic disassembly

Based largely on genetic evidence, we previously proposed that UNC-8 is expressed in DD neurons to promote synaptic remodeling and that the downstream mechanism depends on the elevation of intracellular Ca^2+^ by a positive feedback loop involving UNC-8/DEG/ENaC, UNC-2/VGCC and TAX-6/CaN (Figure 1 B) (Miller-Fleming et al., 2016, 2021; Petersen et al., 2011). Here, we have tested key predictions of this model for UNC-8-dependent synapse elimination (Miller-Fleming et al., 2016) (Figure 1B). First, we have shown that endogenous UNC-8 expression is upregulated at the onset of DD synaptic remodeling and thus could function as a temporal activator of synapse elimination (Figure 1E). Second, UNC-8 localizes to the DD presynaptic zone during remodeling (Figure 1H) and UNC-8 trafficking into the axon depends on UNC-104/KIF1A, the canonical kinesin-3 motor for anterograde transport of presynaptic components (Figure 1F-G) (Hall & Hedgecock, 1991; Otsuka et al., 1991). Third, UNC-8 is required for elevated levels of presynaptic Ca^2+^ in remodeling DD neurons as predicted by previous genetic evidence (Figure 2) (Miller-Fleming, 2016). Fourth, the Ca^2+^-dependent phosphatase TAX-6/CaN is constitutively enriched at presynaptic boutons of GABAergic neurons (Figure 3B-E). Fifth, consistent with the proposed positive-feedback loop that upregulates Ca^2+^ signaling in remodeling boutons (Figure 1B), TAX-6/CaN enhances Ca^2+^ responses during the remodeling window (Figure 3I). Together, these results provide strong support for the hypothesis that UNC-8 elevates intracellular Ca^2+^ in a positive feedback loop involving TAX-6/CaN and that this effect promotes presynaptic disassembly in remodeling DD neurons (Figure 1B)(Miller-Fleming et al., 2016).

### A mechanism of bulk endocytosis drives DD neuron synaptic remodeling

The remodeling pathway that we have identified resembles that of Activity-Dependent Bulk Endocytosis or ADBE. During periods of high synaptic activity, ADBE mediates the recovery of presynaptic membrane for recycling into the synaptic vesicle (SV) pool (Emma L Clayton & Cousin, 2009). For ADBE, elevated intracellular Ca^2+^ activates the phosphatase, Calcineurin/CaN, which, in turn, dephosphorylates a group of conserved proteins (“dephosphins”) that direct endocytosis (Cousin & Robinson, 2001; Guiney et al., 2015). Dephosphorylation of the GTPase, Dynamin, for example, mediates the formation of a Dynamin-Syndapin complex to promote branched-actin polymerization, which drives the formation of bulk endosomes (Figure 4A) (Emma L Clayton & Cousin, 2009). Notably, clathrin is not required for bulk endosome formation but does promote budding at a later step to produce recycled SVs (Emma L Clayton & Cousin, 2009).

We have proposed that TAX-6/CaN acts in a positive feedback-loop with UNC-8 to maintain high Ca^2+^ levels at remodeling synapses (Miller-Fleming et al., 2016) (Figure 1B). Our new results suggest that TAX-6/CaN is also required downstream of UNC-8 to promote an ADBE-like mechanism that dismantles the presynaptic apparatus (Figure 4A-C). Consistent with this idea, we used genetic analysis to confirm that multiple dephosphins (Cousin & Robinson, 2001) promote the elimination of presynaptic domains in remodeling GABAergic neurons (Figure 4D & S4D-E). Strikingly, a phospho-resistant dynamin protein is sufficient to accelerate DD remodeling, as predicted from a model in which CaN-dependent dephosphorylation of dynamin drives endocytic removal of presynaptic domains (Figure 4E-G). In addition, we showed that the F-BAR protein and known ADBE effector, SDPN-1/Syndapin, localizes to remodeling synapses, functions in a common pathway with UNC-8 and mediates UNC-8-dependent elimination of ventral synapses in remodeling GABAergic neurons (Figure 5A-D & S5A-B). Our finding that clathrin is not required for presynaptic disassembly (Figure S4D) is also suggestive of an ADBE-like mechanism since bulk endocytosis is clathrin independent (Emma L Clayton & Cousin, 2009). Finally, we used EM reconstruction to confirm that bulk endosomes, an ultrastructural hallmark of ADBE, are abundantly distributed in remodeling DD neurites and that the UNC-8 protein promotes their formation (Figure 7A-E). Together, these findings are consistent with the hypothesis that an ADBE-like mechanism drives DD synaptic remodeling. In this case, however, bulk endocytosis effectively dismantles the DD presynaptic domain as opposed to the canonical role of ADBE of sustaining presynaptic function (Emma L Clayton & Cousin, 2009).

In addition to the similarities between ADBE and the proposed endocytic mechanism for DD synaptic remodeling, we also identified at least one difference that could be related to the strikingly distinct outcomes of synaptic maintenance vs destruction. During ADBE, Syndapin is proposed to recruit N-WASP to activate the Arp2/3 complex (Alexandros C. Kokotos & Low, 2015). However, N-WASP/WSP-1 protein is dispensable for remodeling GABAergic synapses (Figure S5F). In contrast, the Wave Regulatory Complex (WRC), which also activates Arp2/3 (Machesky et al., 1999), is necessary and functions downstream of UNC-8-dependent synapse elimination (Figures 5F-G and S5F). The observation that WRC, but nor WSP-1, is required for the DD synaptic recycling mechanism is also consistent with the finding that the WRC drives endocytosis in the *C. elegans* intestine and coelomocytes (Patel & Soto, 2013)

### Actin polymerization promotes presynaptic refinement

We used live-imaging to confirm that actin polymerization is elevated in presynaptic domains during the developmental period of DD remodeling and that transient RAB-3 clusters are detected in these regions (Figures 6 and S6). We showed that actin polymerization is reduced in *unc-8* mutants and correlates with fewer transient RAB-3 events and with delayed elimination of stable RAB-3 clusters (Figure 6). The observed actin polymerization defect in *unc-8* mutants is consistent with our finding that the F-BAR protein, SDPN-1/Syndapin, which can recruit nucleators of branched-actin polymerization (Kessels & Qualmann, 2004), functions downstream of UNC-8 (Figure 5A-D). Our finding that actin polymerization is elevated during DD remodeling highlights the need for free barbed ends for Arp2/3-dependent nucleation, as also observed at the leading edge of migrating cells (Falet et al., 2002; Pollard & Borisy, 2003). This requirement could explain an earlier report that the Ca^2+^-sensitive F-actin severing protein, gelsolin, promotes DD synapse elimination (L. Meng et al., 2015). Perhaps gelsolin activity is needed to liberate barbed ends to restructure the actin cytoskeleton for endocytosis and presynaptic disassembly. The additional observation that gelsolin is Ca^2+^-sensitive (Kinosian et al., 1998), also suggests that synaptic Ca^2+^ levels maintained by UNC-8 might promote gelsolin activation.

### Presynaptic proteins recycle from old to new boutons during circuit refinement

We used RNAi knockdown to determine that the recycling endosome component, RAB-11, acts downstream of UNC-8 to remove the SV protein, SNB-1/Synaptobrevin from ventral synapses in remodeling GABAergic neurons (Figure 7F-H). This finding suggested that SV proteins from ventral DD synapses might be recycled for assembly at dorsal synapses. To test this idea, we used a photoconvertible Dendra-2 tag to confirm that the SV-associated protein RAB-3 is recycled to nascent dorsal synapses in remodeling DD neurons (Figures 8 & S8) (Park et al., 2011). Cell-specific RNAi (csRNAi) of *rab-11* in DD neurons, retarded the removal of RAB-3 from ventral synapses as well as impaired its addition to dorsal DD synapses thus confirming that RAB-11 function is required in DD neurons for synaptic recycling (Figure 8E-F). Interestingly, *unc-8* mutants impair dorsal assembly of photoconverted RAB-3 but do not delay its removal from ventral DD synapses (Figure 9), perhaps due to a partially redundant remodeling pathway that acts in parallel to *unc-8* (Miller-Fleming et al., 2021).

Our *in vivo* imaging of RAB-3 and actin dynamics revealed increased actin polymerization as well as mobile RAB-3 particles transiting between ventral DD presynaptic domains during the onset of DD synaptic remodeling (Figure 6). The appearance of transient synaptic vesicles (i.e., mobile GFP::RAB-3 particles) in intersynaptic regions has been previously proposed to drive synaptic vesicle (SV) sharing across *en-passant* boutons to maintain synaptic function in mammalian neurons (Darcy et al., 2006; Herzog et al., 2011; Staras et al., 2010). Moreover, SV sharing between mammalian synapses is also actin-dependent (Chenouard et al., 2020; Darcy et al., 2006; Ratnayaka et al., 2011). Notably, the extent of vesicle sharing declines with the distance between these *en passant* synapses (Staras et al., 2010). Thus, we suggest that a pathway commonly used for local recycling may have been repurposed, in the case of remodeling DD neurons, for recycling of SV components to distal synapses in a mechanism that rewires a developing circuit (White et al., 1978).

In summary, we have shown that a developmentally regulated pathway that dismantles presynaptic domains for reassembly at new locations depends on key elements of a homeostatic mechanism, ADBE, that normally acts to sustain local synaptic function. The widespread occurrence of synaptic refinement across species and the evolutionary conservation of the key components (e.g. DEG/ENaC, Calcineurin, Dynamin, Syndapin, WRC, Arp2/3, RAB11) that drive synaptic remodeling in *C. elegans*, suggest that similar mechanisms may be broadly utilized in the developing brain.

## Star Methods

### Resource Availability

#### Lead contact

Further information and requests for reagents should be directed to and will be fulfilled by the lead contact, David M. Miller, III

#### Materials availability

Key *C. elegans* strains in this study have been deposited at the CGC

#### Data and code availability

## Experimental Model and Subject Details

### Strains and Genetics

Worms were maintained at 20°-23°C using standard techniques (Brenner, 1974). Strains were maintained on NGM plates seeded with *E. coli* (OP-50) unless otherwise stated. The wild type (WT) is N2 and only hermaphrodite worms were used for this study.

#### Generation of unc-8::gfp11×7(syb1624) and tax-6::gfp11×3(sy2783b) animals

Sunybiotech used CRISPR/Cas9 to add 7 copies of GFP11 to the *unc-8* locus and 3 copies of GFP11 to the *tax-6* locus animals at the C-terminus. To visualize the reconstituted GFP signal in DD neurons, complementary GFP1-10 was driven with the P*flp-13* promoter (He et al., 2019).

#### Generation of rab-3(syb2844) [Dendra-2 FLP-ON]) II

Sunybiotech used CRISPR/Cas9 to add a FLP-ON cassette at the N-terminus of the *rab-3* coding sequence (Schwartz & Jorgensen, 2016), which contains the photoconvertible protein Dendra-2. To visualize the endogenous RAB-3 tagged with Dendra-2, we expressed flippase in DD neurons using the *flp-13* promoter (P*flp-13*). Animals were outcrossed three times before they were used for experiments.

## Method Details

### Worm synchronization and staging

Staging was achieved by picking 30-40 L4 larvae onto a fresh plate in the afternoon and leaving at 23C overnight. The next morning, once the animals had become gravid adults, 20-30 worms were picked from original plate onto a fresh plate and allowed to lay eggs for 1 hr at RT. Adults were then removed maintained at 23C to allow the eggs (embryos) to hatch and larvae to develop until selected timepoints for imaging which were defined in as “hours post lay” or “hpl.” Early L4 larvae were identified under DIC (Differential Interference Contrast) imaging in a compound microscope as animals with vulval shapes matching L4.0, L4.1 or L4.2 morphological categories (Mok et al., 2015).

### Molecular Biology

Most plasmids were constructed using InFusion cloning. Plasmid pACC161 (P*flp-13*::dyn-1AAA::SL2::3xNLSTagRFP) was built using Q5 site-directed mutagenesis kit. The pACC156 (P*flp-13*::dyn-1::SL2::3xNLSTagRFP) plasmid was used as a template to change three residues in the phospho-box for a triple-alanine mutant.

### Synapse density analysis

To analyze puncta density along the ventral cord, Maximum Intensity Projections were created from Z-stacks using NIS Elements. All images were subjected to background correction using the Rolling Ball algorithm. Analysis explorer was used to create a mask based on fluorescent thresholding for the GFP::RAB-3 signal. ROIs (Regions Of Interest) were defined for the interval spanning DD2 and DD3 ventral neurites. To calculate puncta density, each object was considered a RAB-3 punctum and the total number was normalized to a 10 μm neurite using the following equation: (# of objects detected in ROI / length of ROI)*10.

To quantify GFP::RAB-3 density in the dorsal nerve cord, ROIs were drawn within the anatomical DD1-DD3 region. Because curvature of the dorsal cord in the XY plane could artificially alter puncta density, all dorsal chord regions used in our analysis had to meet straightness criteria. To this end, the Angle measurement tool in NIS elements was used to measure the angle of intersection between two lines drawn tangent to the regions of the dorsal cord immediately anterior and posterior to the apex of the curve in question. Regions with an intersection measurement of less than 160° (where 180° represents a perfectly straight region and 90° represents a region curved at a right angle) were excluded from analysis. ROIs were then drawn along the longest possible region of the dorsal cord that adhered to these straightness criteria. If the regions anterior and posterior to a curve met straightness criteria, then two ROIs were drawn for that image. Off target fluorescent features (autofluorescence) and commissures were excluded from ROIs. ROI lengths were measured using the Distance Measurement: Polyline tool in NIS Elements and were recorded in μm. The Automated Measurement Results tab in NIS Elements was used to record the number of objects (puncta) in each ROI, and puncta density within the ROI was then calculated as described above. To designate a fluorescent signal as a punctum, a mask was generated in NIS Elements that labeled as puncta all fluorescent features in an image that met size, shape and intensity criteria set by the experimenter. Once generated, the same masking criteria were used for all treatment and control images included in a given experiment.

### Counting UNC-8::GFP11_x7_ puncta

Worms were synchronized (see above) to track endogenous UNC-8 NATF GFP puncta before and during the remodeling window: early L1 (16hpl), late L1 (14hpl) and L2 stage (30hpl). Z-stacks were collected using a Nikon A1R laser scanning confocal microscope (40x, 1.4 N.A.). Worms were immobilized in a pool of 3μL of 100mM muscimol (TOCRIS biosciences #0289) + 7μL 0.05um polybeads (2.5% solids w/v, Polysciences, Inc. #15913-10) on a 10% agarose pad. Maximum Intensity projections were created for each image using NIS Elements. The experimenter was blinded to developmental time to manually score each GFP punctum. UNC-8::GFP puncta density was calculated by normalizing the number of puncta to a 10 μm neurite.

### Presynaptic TAX-6 enrichment

Worms were synchronized (see above) to track endogenous TAX-6::GFP11_x3_ NATF fluorescence along the ventral and dorsal cords before at the L1 stage (16hpl), and after remodeling at the L4 stage. Z-stack images were acquired using a Nikon A1R laser scanning confocal microscope (40x, 1.4 N.A). Maximum intensity projections were created using FIJI and a 3-pixel wide line scan was drawn on the ventral and dorsal cord of each animal to determine average fluorescence. The relative dorsal (D) and ventral fluorescence (V) was normalized to the total fluorescence (D+V) for each DD neuron (DD1-DD4).

To detect endogenous TAX-6::GFP and mRuby::CLA-1s, Z-stacks were acquired using Nyquist Acquisition and small step size (0.2 μm) on a Nikon A1R confocal microscope. Z-stacks were subjected to 3D-deconvolution in NIS Elements and single plane images were used for line scans.

### GCaMP6s imaging in remodeling DD axons

To detect GCaMP6s fluctuations at remodeling boutons (DD2-DD4), NC3569 animals were synchronized (26hpl) on an OP-50-seeded plate with freshly added ATR or carrier (EtOH, no ATR). Image acquisition was performed on a Nikon TiE microscope equipped with a Yokogawa CSU-X1 spinning disk head, Andor DU-897 EMCCD camera, high-speed MCL piezo and 100X/1.49 Apo TIRF oil objective lens. Synchronized NC3569 worms at 26hpl (See synchronization method above) were immobilized in a pool of 3μL of 100mM muscimol (TOCRIS biosciences #0289) + 7μL 0.05um polybeads (2.5% solids w/v, Polysciences, Inc. #15913-10) on a 10% agarose pad.

To identify baseline GCaMP fluctuations, images 4-10fps (frames per second) were collected with 488 nm excitation in a single plane using a Perfect Focus System for 1 minute. The ventral processes of DD2-DD4 were imaged for this experiment. To detect evoked GCaMP changes, triggered acquisition was used to excite GCaMP with 488 nm and activate Chrimson with 561nm laser. Single plane movies were collected at 3.8 fps for 1 minute using emission filters 525nm (+/-25nm) and 646 (+/-66nm). The sample was illuminated with a 561nm laser at 5 sec intervals (e.g., every 20^th^ frame) to activate Chrimson expressed in cholinergic DA motor neurons of late L1 animals (26hpl) (*Punc-4:*:ceChrimson::SL2::3xNLS::GFP) while maintaining constant illumination with a 488 nm laser to detect GCamP6s signals. This acquisition paradigm was applied for 10 cycles (1 minute). Only neurites that were stable (not moving or sinking) for ≥ 2 cycles of acquisition were selected for further processing. For quantifying GCaMP6s fluorescence, ROIs were drawn on oblong, bouton-like structures near the base of the commissure and on a nearby region to capture background fluorescence. The same ROIs (bouton and background) were used to detect spontaneous and evoked GCaMP changes in videos from the same DD neuron. Mean fluorescence intensity of each ROI for each frame was exported into Excel for analysis. Background was subtracted from each frame and measurements were normalized to three time-points before Chrismon activation for the first acquisition cycle (F_0_). At each time-point, F_0_ was subtracted from the measured GCaMP fluorescence and then divided by F_0_ to determine ΔF/F_0._ Then, fluorescent values before and after each event of Chrimson activation were matched for paired comparisons.

### Cell-specific RNAi

We used In-Fusion cloning to create two plasmids that express complementary strands of mRNA for each cell-specific RNAi (csRNAi) experiment. To confirm expression, each plasmid contains a transplicing sequence (SL2) followed by a fluorescent protein (cytosolic of nuclear-localized GFP and mCherry) downstream of the sense or antisense transcript. For example, for csRNAi knockdown of *arx-5, dyn-1, rab-11* and *rab-5* in DD neurons, we drove expression of one strand in DD neurons with the *flp-13* promoter and the opposite strand in DD + VD neurons with the *ttr-39* promoter (See Table 1). In this arrangement, expression of both the sense and anti-sense sequences is limited to DD neurons, thus resulting in a DD-specific knockdown of each targeted gene. csRNAi templates for each target gene were generated by PCR from genomic DNA with primers that excluded the ATG start codon as follows: 1210 bp of *p21/arx-5*, 1686 bp or *dyn-1*, 1174 bp of *rab-11*.*1* and 1275bp of *rab-5*.

In this experimental design, csRNAi-treated DD neurons are marked by expression of both fluorescent markers (mCherry + GFP) whereas control DD neurons are unlabeled. To determine if remodeling of SNB-1::GFP (*juIs137*) was perturbed upon knockdown, ImageJ software was used to generate maximum intensity projections. All fluorescence intensity plots were created by drawing a linescan through the ventral and dorsal nerve cords of each worm and averaging the fluorescence intensity values across the anterior region of the DD neurons scored. Intensity of SNB-1::GFP was normalized to the total dorsal and ventral fluorescence.

### Feeding RNAi

Clones from the RNAi feeding library (Source BioScience) were used in this study (Lisa Timmons & Andrew Fire, 1998)(Kamath et al., 2003). RNAi plates were produced as previously described (Earls et al., 2010; Petersen et al., 2011). First, 15 mL culture tubes containing 2 ml LB broth with 10 μl (10mg/mL) ampicillin were inoculated with individual bacterial colonies and grown in a 37° C shaker for 12-16 hr. Control RNAi cultures were inoculated with bacterial colonies containing the cloning vector but no insert (“empty vector” control) for each feeding RNAi experiment. 250 μL of overnight culture was added to a mixture of 12.5 ml LB broth + 62.5 μl ampicillin in a 50 mL conical tube. The new culture was then returned to the 37 C shaker until reaching log phase (OD > 0.8). The culture was then induced by adding 12.5 mL of LB broth, 62.5 μL ampicillin, and 100 μL 1M IPTG. After 3.5-4 hours incubating in 37° C shaker, the culture was spun down using a tabletop centrifuge for 6 min at 3900 rpm to pellet the bacteria. The supernatant was discarded, and the pellet resuspended in 1mL M9 mixed with 8μL IPTG. The bacterial mixture was dispensed onto 4 × 60 mm NGM plates (250 μl each), and the plates were left to dry overnight in the horizontal laminar hood and then stored at 4° C for up to a week. To set-up each RNAi experiment, 3-5 L4 larvae (NC1852) were placed on each RNAi plate and maintained at 20°C. Four days later, F1 progeny was imaged as L4 animals. To count the number of puncta, see below.

### Identifying remodeling genes

To screen candidate genes for a role in GABAergic synapse remodeling, we scored SNB-1::GFP puncta from the *juIs1* [p*unc-25*::SNB-1::GFP; *lin-15*] transgene (Hallam & Jin, 1998) in *unc-55; eri-1* mutants sensitive to feeding RNAi as previously described (Petersen et al., 2011). We compared animals treated with each candidate gene RNA feeding strain vs the blank RNAi (empty vector). Briefly, animals were anesthetized with 0.1% tricaine/tetramisole, mounted on a 2% agarose pad, and imaged with an inverted microscope (Zeiss Axiovert) using Micro Manager software and a 63X oil objective. Ventral puncta between VD3 to VD11 were counted following the mid-L4 stage for both RNAi control and RNAi-treated animals. Data were pooled at least from 2 separate experiments and the examiner was blinded to the treatment. One-Way ANOVA was performed with post hoc correction to compare each RNAi treatment against the control. Similarly, the scorer was blinded to genotype to test if Synaptojanin/*unc-26* promotes synapse remodeling of SYD-2::GFP puncta from *hpIs3[punc-25::SYD-2::GFP; lin-15*] transgene (Yeh et al., 2005) in wild-type, *unc-55* and *unc-55; unc-26* double mutants.

### *dyn-1*(OE) analysis

Synchronized animals (28hpl) (28 hours post laying) expressed either wild-type (WT) or phospho-resistant versions of *dyn-1* in DD neurons under the P*flp-13* promoter in tandem with a nuclear-localized TagRFP protein to recognize cells expressing the array. Z-stacks were collected in a Nikon A1R laser scanning confocal spanning the depth (∼ 3 microns) of the ventral nerve cord. Z-stacks were used to generate MaxIPs in NIS Elements and Analysis Explorer, to define a binary mask based on fluorescence and size to recognize cell soma that expressed nuclear TagRFP. DD1-DD3 neurons were categorized as Control (no red) or *dyn-1*(OE) (red nucleus) based on nuclear expression of TagRFP. An additional mask was defined in the GFP channel for ventral SNB-1::GFP *(juIs137)* puncta based on fluorescence and circularity. Each recognized punctum on the anterior process of all DD1-DD3 neurons was counted. Then, puncta counts for each DD dendrite was normalized to the length of the counted region to calculate SNB-1::GFP density as puncta/10 μm.

### VCA-1(OE) analysis

Synchronized animals (26hpl) expressed the VCA domain of the Wave Regulatory Complex in tandem with cytosolic mCherry with the P*ttr-39* promoter to mark GABAergic cells carrying the array. Z-stacks spanning the depth of both nerve cords were collected in a Nikon A1R laser scanning confocal microscope. Z-stacks were used to generate MaxIPs in FIJI and the segmented line tool (3-pixel wide) was used to draw ROIs on top of the ventral and dorsal nerve cords. mCherry-positive DD cells carry a VCA(OE) transgenic array. All fluorescence intensity plots were created by drawing a segmented line through the ventral and dorsal nerve cords of each worm and averaging the fluorescence intensity values across the anterior region of the DD neurons scored. Intensity of SNB-1::GFP (*juIs137*) was normalized to the total dorsal and ventral fluorescence.

### UNC-8(OE) analysis in RNAi-treated animals

FIJI was used to quantify presynaptic markers in VD neurons for effects of UNC-8(OE) (over-expression). Z-stacks were collected for the full length of the ventral nerve cord (VD3 – VD11). mCherry-positive VD cells carry an UNC-8 cDNA transgenic array (Miller-Fleming et al., 2016). Because UNC-8(OE) transgenic arrays are mosaic with expression limited to a random subset of VD neurons in each animal, data were collected from VD neurons (VD3-VD11) carrying the UNC-8 cDNA (mCherry-positive) vs an adjacent control VD neuron that does not carry the array. Neighboring mCherry-positive [e.g., UNC-8(OE)] and mCherry-negative [control] VD neurons were compared to quantify differences in the fluorescence signal for SNB-1::GFP due to UNC-8 over-expression with RNAi knockdown of either *tax-6, sdpn-1, wve-1* or *rab-11*. Intensity values were obtained from line scans anterior to the VD cell bodies of interest. Background fluorescence was obtained from a line scan of an adjacent region inside the animal and subtracted from the VD line scans.

### Microscopy

#### Laser Scanning Confocal Microscopy

Unless otherwise indicated for specific experiments, Larval or young adult animals were immobilized on 2-10% agarose pads with 15mM levamisole, 0.05% tricaine as previously described (Smith et al., 2010). Z-stacks were acquired with a Nikon confocal A1R using Apo Fluor 40X/1.3 and 60X/1.4 N.A. oil objectives.

#### In-vivo actin dynamics at remodeling synapses

For measurements of actin dynamics at synaptic regions in DD neurons (Figures 6 and S6), endogenous GFP::RAB-3 was used to mark synaptic vesicle clusters and LifeAct::mCherry (Riedl et al., 2008) to visualize actin dynamics. Live-imaging was performed in a Nikon TiE microscope equipped with a Yokogawa CSU-X1 spinning disk head, Andor DU-897 EMCCD camera or Photometrics Prime 95B sCMOS camera, high-speed piezo stage motor, a perfect focus system and a 100X/1.49 Apo TIRF oil objective lens. Synchronized animals at 16-18hpl (early L1) or 26-28hpl (late L1) were mounted on 10% agarose pads and immobilized in a pool of 3μL of 100 mM muscimol (TOCRIS biosciences #0289) + 7μL 0.05um polybeads (2.5% solids w/v, Polysciences, Inc. #15913-10). Single-plane snapshots were collected at 1 fps using triggered acquisition. Movies were submitted to 2D-deconvolution on NIS-Elements using the Automatic algorithm and aligned with the NIS Elements alignment tool when necessary. To analyze GFP::RAB-3 dynamics, videos were limited to 60-150 seconds to reduce photobleaching and the GFP signal was binarized with manual thresholding to enhance local contrast. The resultant binarized video was then submitted for particle tracking to detect transient GFP::RAB-3 movement. The total number of transient events was normalized to events per minute per 10 μm neurite. Kymographs to visualize GFP::RAB-3 and actin dynamics (LifeAct::mCherry) were generated in NIS Elements. At sites of GFP::RAB-3 puncta, ROIs were defined in the 561nm channel to track LifeAct::mCherry transients (e.g, actin transients) in presynaptic regions. LifeAct::mCherry fluorescence throughout time were plotted and normalized with the first fluorescence value set to zero. GFP::RAB-3 density (puncta/10 micron) for each genotype was determined from the first frame of each video and submitted for 2D deconvolution with NIS elements. The line scan function was then used to define fluorescent GFP::RAB-3 peaks. Any peak at > 300AU fluorescent units was considered a GFP::RAB-3 punctum.

#### In-vivo dynamics of endogenous GFP::RAB-3 and TagRFP::RAB-11 during remodeling

We built a strain to follow endogenous GFP::RAB-3 and TagRFP::RAB-11 dynamics (Figure S7D-F) during the remodeling window 27-30 hpl. Animals were placed on 10% agarose pads and immobilized using a combination of 3μL of 100mM muscimol (TOCRIS biosciences #0289) and 7μL 0.05um polybeads (2.5% solids w/v, Polysciences, Inc. #15913-10). Single plane movies were captured at 0.5fps (frames per second) on a Nikon TiE microscope with a Yokogawa CSU-X1 spinning disk head, Photometrics Prime 95B sCMOS camera, a high-speed MCL Piezo stage, a Perfect Focus system and a 100X/1.49 Apo TIRF oil objective lens. Movies were 2D-deconvolved using the automatic algorithm in NIS Elements and kymographs were created in FIJI with the Multi Kymograph tool. Kymographs were then exported and submitted to Kymobutler (Jakobs et al., 2019) for identification of particle movement (https://www.wolframcloud.com/objects/deepmirror/Projects/KymoButler/KymoButlerForm).

#### Tracking of endogenous Dendra-2::RAB-3 during remodeling

This experiment consists of four steps: (1) worm synchronization, (2) photoconversion and collection of baseline data before remodeling, (3) worm recovery, and (4) analysis of photoconverted signal after remodeling. Larvae expressing endogenous Dendra-2::RAB-3 in DD neurons, were synchronized by allowing young adults to lay eggs for 1 hr (see above). Plates were maintained at 23 C. L1 animals at 18 hpl were photoconverted on a Nikon TiE microscope with a Yokogawa CSU-X1 spinning disk head using a Mini Scanner equipped with a 405 nm, 100mW photo stimulation laser. Briefly, single worms were mounted on a 10% agarose pad with 2μL of anesthetic [from mixture of 3μL of 100mM muscimol (TOCRIS biosciences #0289) and 7μL 0.05um polybeads (2.5% solids w/v, Polysciences, Inc. #15913-10)]. Wax-paraplast 50/50 v/v combination was applied to each corner of the coverslip to secure it. Using NIS elements, we defined a targeted ROI limited to ventral Dendra-2::RAB-3 synaptic puncta visualized in the 488 nm channel. Dendra-2::RAB-3 was photoconverted with 30μs dwell time with 0.5% power of the 405 nm laser. We collected a Z-stack using 488nm and 561nm lasers before and after Dendra-2 photoconversion. In most cases, we photoconverted Dendra-2::RAB-3 puncta in the anterior ventral neurites of either DD2, DD3 or DD4.

To recover worms, we added 90μL of M9 to the space between the coverslip and the agarose pad and cut the sealant from each corner using a razor blade. The coverslip and agarose pad were rinsed with the addition of M9 buffer and allowed to drip on top of an NGM 60mm plate. Typically, two rounds of 90μL M9 was used to rinse the coverslip and five rounds for the agarose pad. After a few minutes, when the NGM plates had dried, each plate was placed in a 23C incubator. Each treated animal was maintained on a separate NGM plate to facilitate matching imaging data obtained before and after remodeling. Twenty hours after photoconversion (38 hpl), we mounted each worm on a 10% agarose pad using 3uL of the anesthetic mixture (see above) and collected a Z-stack using 488nm and 561nm lasers to track the Dendra-2::RAB-3 signal.

For analysis, we first used NIS Elements to 3D-deconvolve all Z-stacks and paired the images for each individual worm before (18hpl) and after remodeling (38 hpl). After deconvolution, we imported all images to FIJI and created maximum intensity projections. We drew a 3-pixel wide line scan on top of the Dendra-2::RAB-3 signal and exported the traces into Prism from each fluorescent channel (488 and 561nm) and time-point. Traces from neighboring regions were used to define background fluorescence. We used Prism to determine the area under the curve for line scans of the ROI (Dendra2::RAB-3 puncta) as well as neighboring regions (background). We subtracted the background area and plotted the relative photoconverted signal (vs 18hpl) in the dorsal and ventral side at 38 hpl.

### Electron Microscopy

20 young adults were placed on an OP-50 seeded NGM plate, allowed to lay eggs for one hour and then removed. Timing for hours post laying (hpl) was started halfway through the hour. Cultures were maintained at 23 C. We monitored the endogenous GFP::RAB-3 marker to select late-L1 stage larvae at 27hpl during the remodeling period for high-pressure freeze (HPF) fixation.

Strains were prepared using HPF fixation as previously described (Liu et al., 2021). Briefly, twenty to thirty staged L1 larvae at 27 hpl were placed in specimen chambers filled with *Escherichia coli* and frozen at −180°C, using liquid nitrogen under high pressure (Leica HPM 100). Samples then underwent freeze substitution (Reichert AFS, Leica, Oberkochen, Germany) using the following program: Samples were held at −90°C for 107 hours with 0.1% tannic acid and 2% OsO_4_ in anhydrous acetone, then incrementally warmed at a rate of 5°C/hour to −20°C, and kept at −20°C for 14 hours, before increasing temperature by 10°C/hour to 20°C. After fixation, samples were infiltrated with 50% Epon/acetone for 4 hours, 90% Epon/acetone for 18 hours, and 100% Epon for 5 hours. Finally, samples were embedded in Epon and incubated for 48 hours at 65°C (Liu et al., 2021). Thin (70 nm) serial sections were acquired using an Ultracut 6 (Leica) and collected on formvar-covered, carbon-coated copper grids (EMS, FCF2010-Cu). Sections were post-stained with 2.5% aqueous uranyl acetate for 4 minutes, followed by Reynolds lead citrate for 2 minutes (Liu et al., 2021). Images were obtained using a JEOL JEM-1400F transmission electron microscope, operating at 80 kV. Micrographs were acquired using an AMT NanoSprint1200-S CMOS or BioSprint 12M-B CCD Camera and AMT software (Version 7.01). The DD2 neuron was identified based on measurements taken from 27 hpl larvae with GFP::RAB-3 expressed in DD neurons, and determined using distance from the isthmus of the pharynx in conjunction with established GABA neuron synaptic morphology (White et al., 1986)(Miller-Fleming et al., 2016, 2021).

Serial sections (control = 396; *unc-8* = 349) containing the anteriorly-directed ventral process of the DD2 neuron were manually aligned using the TrakEM2 package of NIH FIJI/ImageJ software. The aligned stack was imported into 3Dmod/IMOD for segmentation and reconstruction of the sections into a 3D model of DD2. Quantitative data were extracted from the 3D model using the *imodinfo* module of IMOD. Values were imported to Prism (GraphPad) for statistical analysis using Fisher’s exact test or t-test. Videos of the model were created using the movie/montage module in IMOD to create TIFF frames and imported into FIJI/ImageJ to create the AVI movie file.

### Statistical Analysis

First, we used the Shapiro-Wilk test to determine if a sample is normally distributed. For comparisons between 2 normally distributed groups, Student’s T-test was used and p<0.05 was considered significant. ANOVA was used for comparisons of > 2 samples and for normally distributed data followed by Dunnett’s multiple-comparison test. Standard Deviations between two samples were compared using an F-test and p<0.05 was considered significant. If the samples were not normally distributed, we used a Mann-Whitney test to compare two groups and a Kruskal Wallis test to compare three or more groups. The specific test and N for each experiment are listed in figure legends. For Fisher’s Exact test in Figure 7C, sections with EVs: WT = 133; *unc-8* = 55 and sections without EVs: WT = 263; *unc-8* = 294. For Fisher’s Exact test in Figure 7SD, sections with dense projections: WT = 21; *unc-8* = 31 and sections without dense projections: WT = 375; *unc-8* = 318.

## Key Resources Table

| REAGENT or RESOURCE | SOURCE | IDENTIFIER |
| --- | --- | --- |
| Bacterial and virus strains |  |  |
| *E. coli* OP50-1 | Caenorhabditis Genetics Center (CGC) | Wormbase ID: OP50-1 |
| Biological samples |  |  |
| Chemicals, peptides, and recombinant proteins |  |  |
| All-trans retinal (ATR) | Sigma-Aldrich | R2500 |
| Critical commercial assays |  |  |
| Deposited data |  |  |
| Experimental models: cell lines |  |  |
| Experimental models: *C. elegans* strains |  |  |
| See Supplemental Table 1 |  |  |
| Oligonucleotides |  |  |
| Recombinant DNA |  |  |
| *punc-4*::Chrimson::SL2::3xNLSGFP | Cuentas-Condori et al., 2019 | pACC92 |
| *pflp-13*::LifeAct::mCherry | Cuentas-Condori et al., 2019 | pACC12 |
| *pttr-39*::VCA::SL2::mCherry | This work | pACC42 |
| *pflp-13*::*arx-5* sense::SL2::mCherry | This work | pACC50 |
| *pflp-13*::*arx-5* antisense::SL2::3xNLS::GFP | This work | pACC51 |
| *pflp-13*::*sdpn-1b*::mCherry | This work | pACC57 |
| *pflp-13*::GCaMP6s::SL2::mCherry | Cuentas-Condori et al., 2019 | pACC83 |
| *pflp-13*::2xNLS::FLP_D5 | This work | pACC98 |
| *pflp-13*::dyn1F::SL2::mCherry | This work | pACC106 |
| *pflp-13*::dyn1R::SL2::3xNLS::GFP | This work | pACC107 |
| *pflp-13::mRuby::CLA-1s* | This work | pACC123 |
| *pflp-13::2xNLS::FLP_D5::SL2::LifeAct::mCherry* | This work | pACC141 |
| *pflp-13::RAB-11.1F::SL2::3xNLSGFP* | This work | pACC150 |
| *pttr-39::RAB-11R::SL2::mCherry* | This work | pACC151 |
| *pflp-13::dyn-1::SL2::2xNLS::TagRFP* | This work | pACC156 |
| *pflp-13::TagRFP::RAB-11.1cDNA* | This work | pACC157 |
| *pttr-39::RAB-11.1csRNAi_R::SL2::2xNLSTagRFP* | This work | pACC158 |
| *pflp-13::dyn-1-AAA::SL2::2xNLSTagRFP* | This work | pACC161 |
| *pmyo-2::RFP* | This work | pSH4 |
| *pstr-1::GFP* | This work | pSH21 |
| *pflp-13::GFP1-10* | He et al., 2019 | pSH87 |
| Software and algorithms |  |  |
| NIS Elements | Nikon | https://www.microscope.healthcare.nikon.com/products/software/nis-elements |
| ImageJ 1.8.0 | NIH | https://imagej.nih.gov/ij/download.html |
| Other |  |  |

**Table 2.**
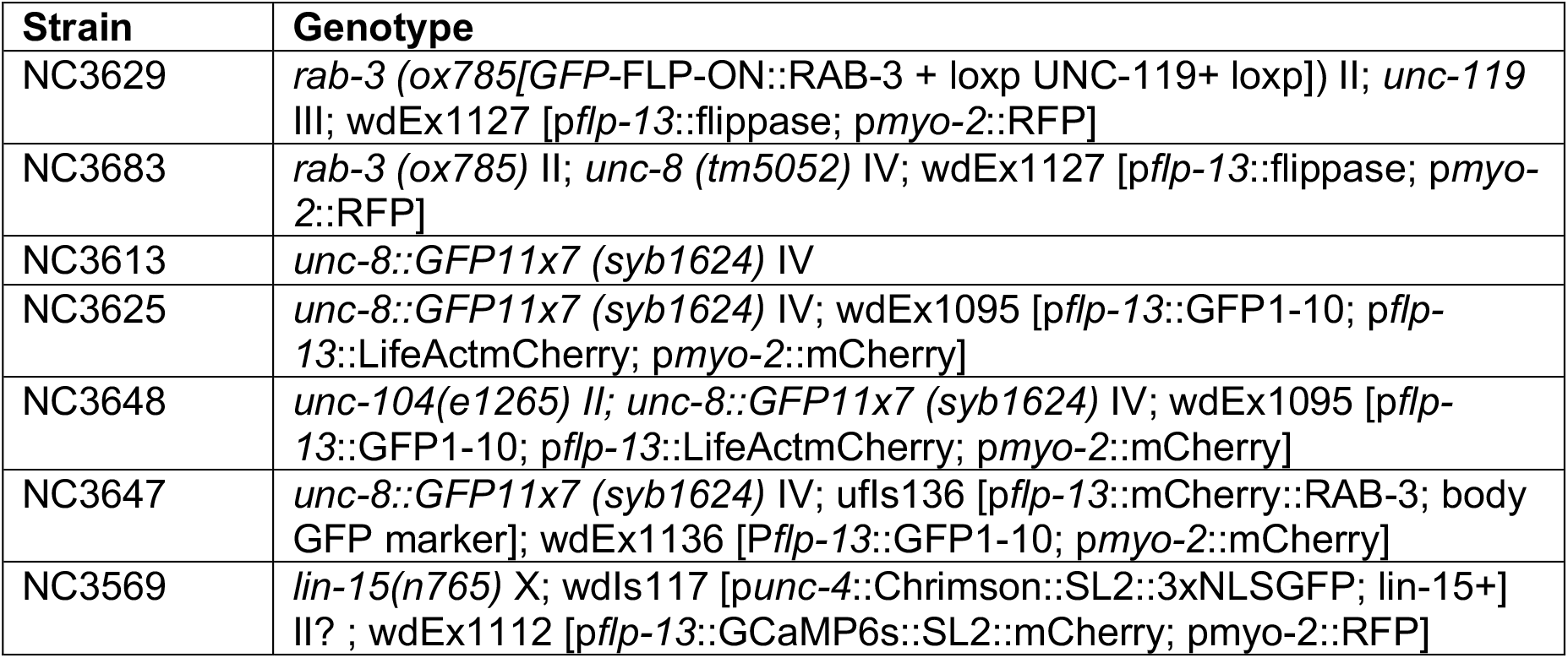
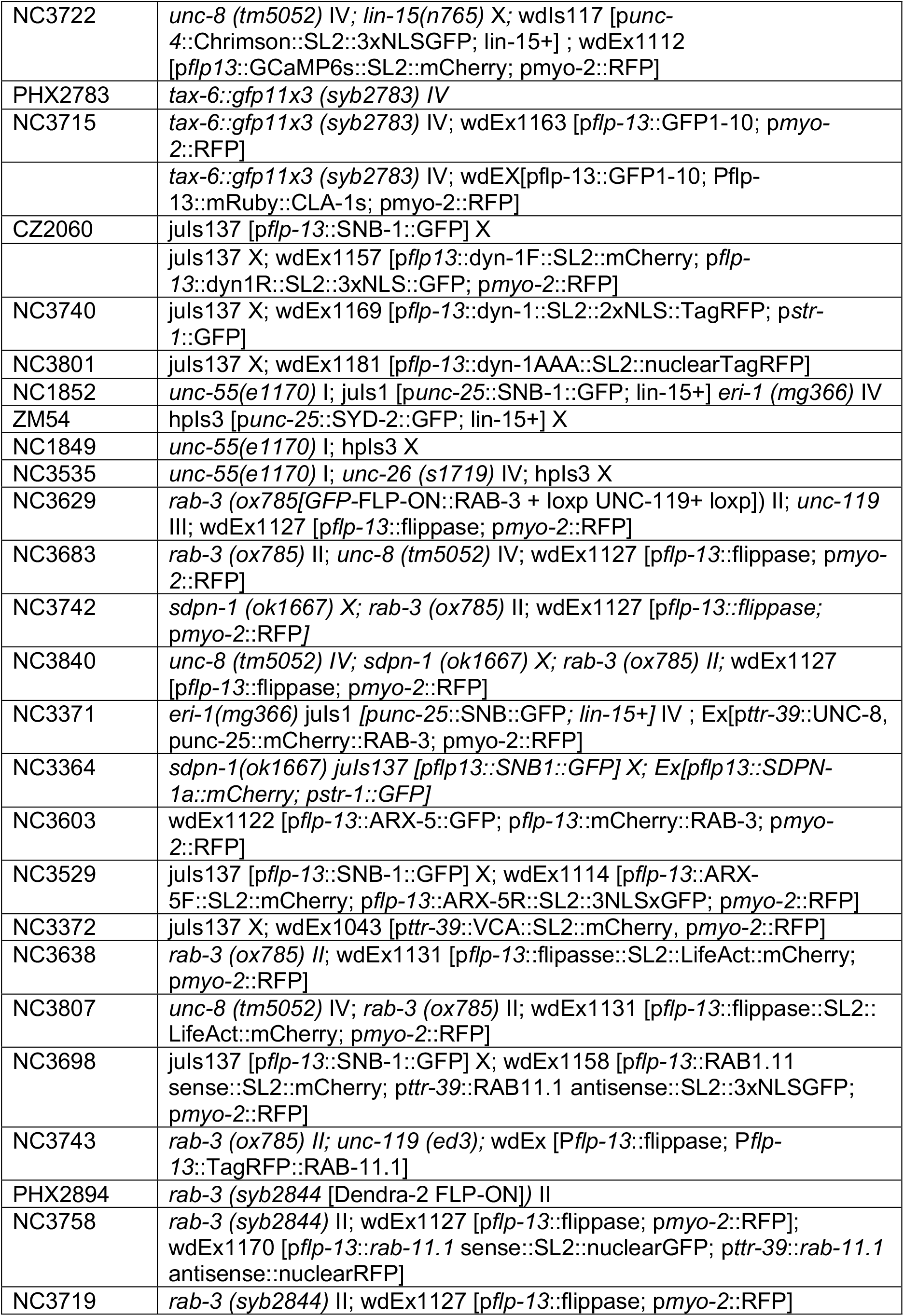
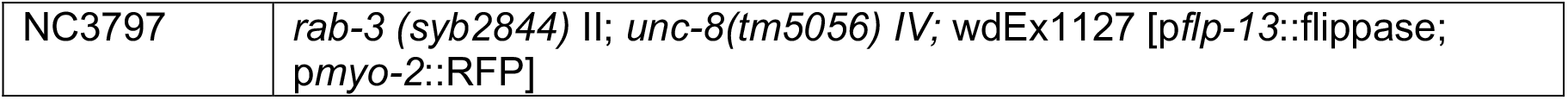
List of strains used in this chapter

## Acknowledgments

We thank Y. Jin, K. Shen and E. Jorgensen for reagents, B. Mulcahy and M. Zhen for sharing unpublished results and B. Millis, N. Grega-Larson and S. Percival for help with calcium and actin imaging. Funded by NIH (R01NS113559) to DMM, AHA (18PRE33960581) to ACC and Vanderbilt Undergraduate Summer Research Program to SC. Experiments were performed in the Cell Imaging Shared Resource which is supported by NIH grant DK020593. Some strains were provided by the CGC, which is funded by NIH Office of Research Infrastructure Programs (P40 OD010440).

## Author Contributions

Conception and design: ACC, KG, MK, JR, DMM; Generation of transgenic lines: ACC; Confocal imaging; ACC, SC, JT, LF; Super-resolution imaging: ACC; Calcium imaging: ACC; electron microscopy, 3D reconstruction and movie: MK; EM analysis: KG, MK, JR; Drafted manuscript: ACC, DMM. Critically revised manuscript: ACC, SC, KG, MK, JT, LF, JR, DMM.

## Declaration of Interests

The authors declare no competing interests.

## Notes

### Competing Interest Statement

The authors have declared no competing interest.

### Summary of Updates

includes videos S1-S9 and legends

