## Supplemental Figures for "The Epithelial Na^+^ Channel UNC-8 promotes an endocytic mechanism that recycles presynaptic components from old to new boutons in remodeling neurons"

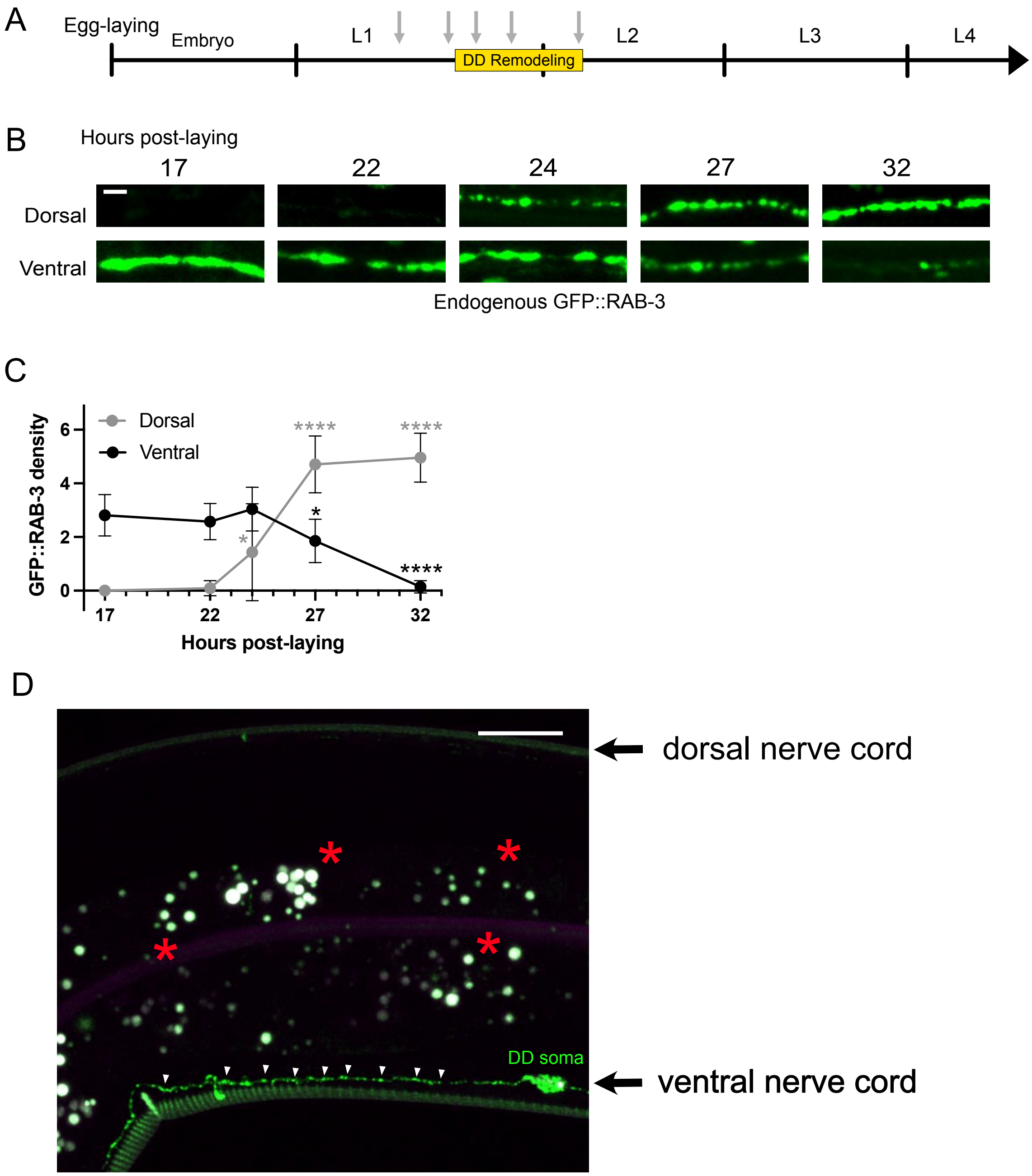
A
Egg-laying
Embryo
L1
DD Remodeling
L2
L3
L4
B
Hours post-laying
17
22
24
27
32
Dorsal
Ventral
Endogenous GFP::RAB-3
C
Dorsal
Ventral
GFP::RAB-3 density
Hours post-laying
D
dorsal nerve cord
DD soma
ventral nerve cord

**Figure 1 – Supplement. Timeline for RAB-3 remodeling in developing DD neurons**

**A.** Timeline depicts development from the embryo through larval stages (L1, L2, L3, L4). DD remodeling (yellow box) spans the transition from the late L1 to early the L2 larval stages at 23C. Arrows (grey) denote five time points,17 to 32 hours post-laying (hpl), evaluated by live imaging of GFP::RAB-3.

**B.** Snapshots of endogenous GFP::RAB-3 in DD neurons at five timepoints encompassing the remodeling period (17-32 hpl). Scale bar = 2 μm.

**C.** GFP::RAB-3 density progressively decreases on the ventral side (black): at 17 hpl (2.81 ± 0.8, n=21 ), 21 hpl ( 2.58 ± 0.7, n=51 ), 24 hpl ( 3.04 ± 0.8, n=22 ), 27 hpl ( 1.86 ± 0.8, n=22 ) and 32 hpl ( 0.14 ± 0.3, n=14 ) as GFP::RAB-3 density increases on the dorsal side (gray): 17 hpl ( 0.0 ± 0, n=12 ), 22 hpl ( 0.09 ± 0.3, n=28 ), 24 hpl ( 1.44 ± 1.8, n=21 ), 27 hpl ( 4.71 ± 1.1, n=15 ) and 32 hpl ( 4.96 ± 0.9, n=14 ). Data are mean ± SD. Kruskal-Wallis test. * $p < 0.05$ and **** $p < 0.0001$.

**D.** Representative image of endogenous UNC-8::GFP (arrowheads) labeled with the NATF strategy localized to a ventral DD dendrite at the L4 larval stage. Asterisks denote gut autofluorescence. Scale bar = 10 μm.

#### Non-ATR controls

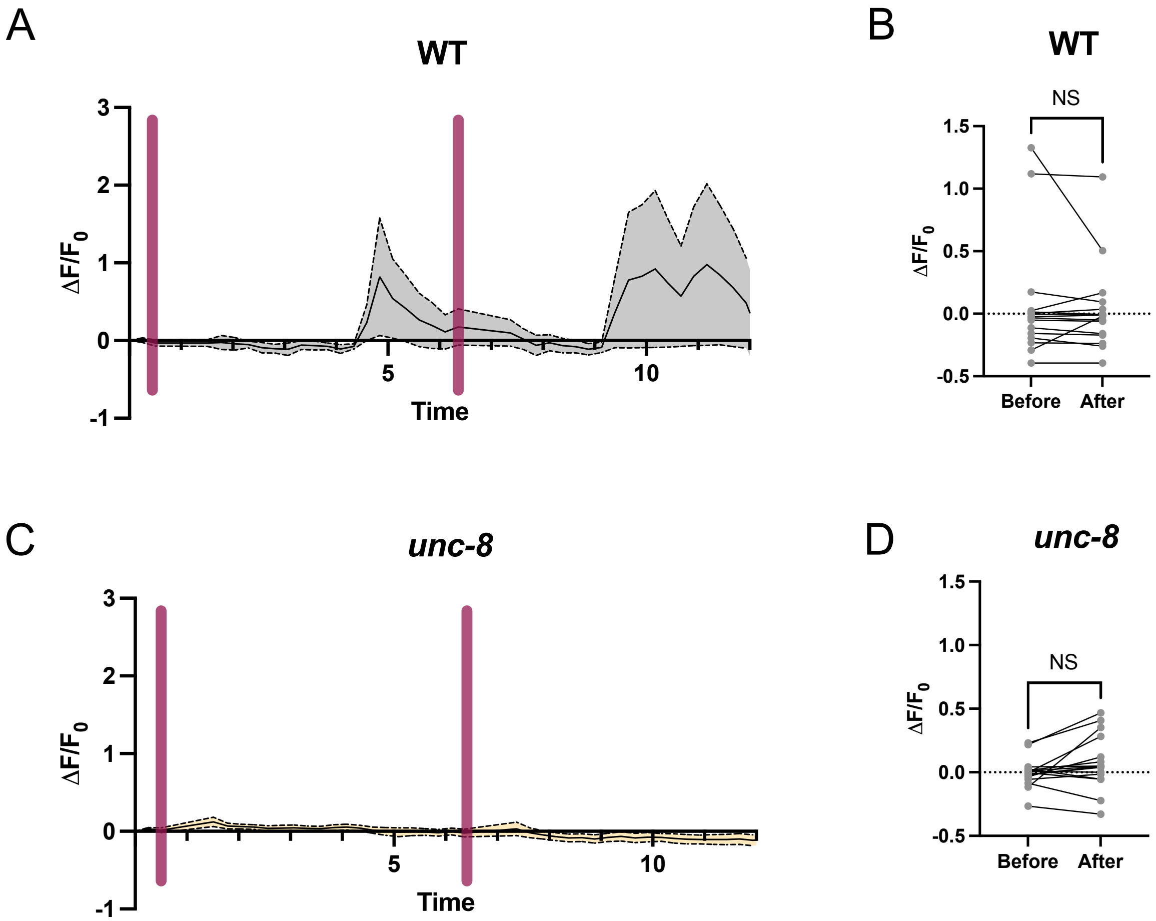


**Figure 2 – Supplement. Non-ATR controls for GCaMP signals in wild type and *unc-8* DD neurons**

In the absence of ATR, Chrimson activation (magenta bars) in DA neurons fails to elevate GCaMP fluorescence in either (**A-B**) wild type (WT) (baseline = 0.07 ± 0.5, n=16; after activation = 0.03 ± 0.4, n=16), p = 0.487 or (**C-D**) *unc-8* mutants (baseline = -0.002 ± 0.1, n=16, after activation = 0.07 ± 0.2, n=16), p = 0.0604. Paired t-test, N.S., Not Significant.

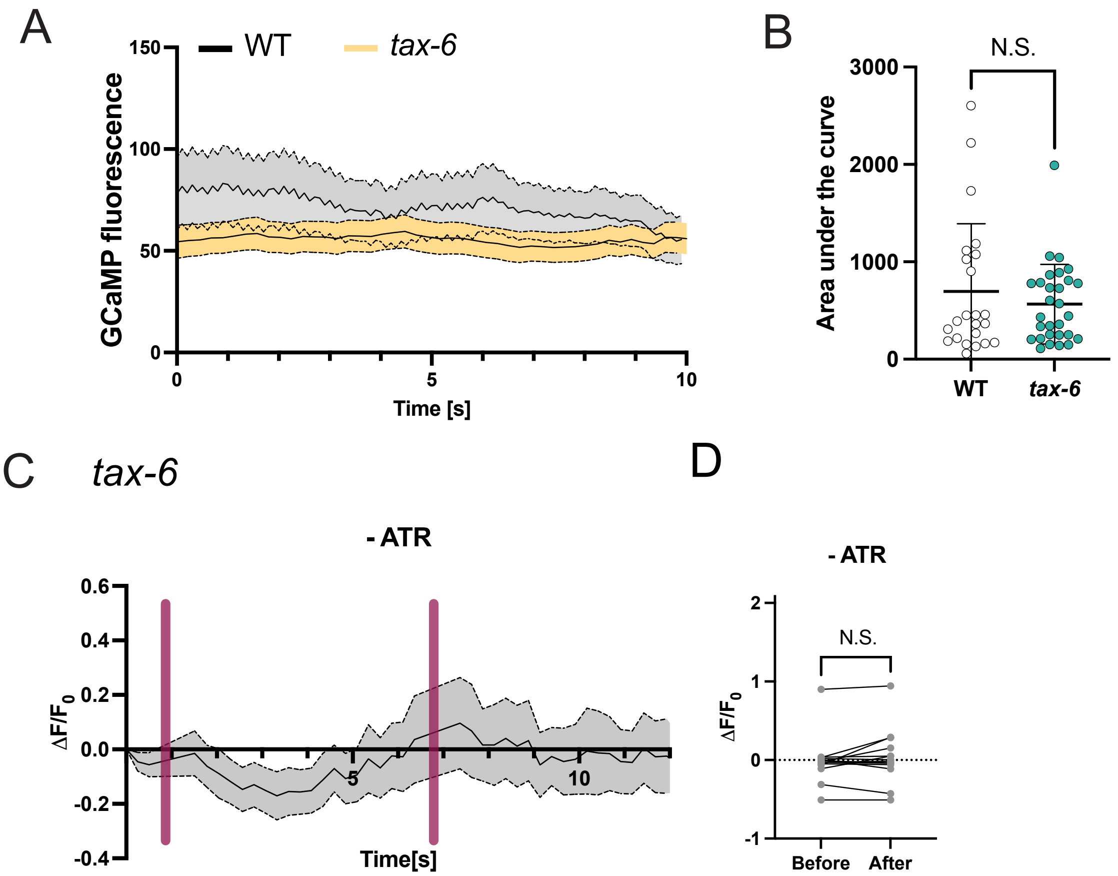


**Figure 3 -Supplement. Spontaneous $Ca^{2+}$ transients in wild-type vs *tax-6* mutant DD neurons**

**A-B.** The magnitudes (area under the curve) of spontaneous $Ca^{2+}$ fluctuations in wild type (WT) (695 ± 145, n=23) (see Figure 2C) vs *tax-6* mutants (566 ± 408, n=29) are not significantly different. Mann-Whitney test, N.S., Not Significant, p = 0.7806. Data for WT also shown in Figure 2.5B.

**C-D. (C)** In the absence of ATR (-ATR), DA activation (magenta bar) does not elevate GCaMP fluorescence in *tax-6* mutant animals (magenta). **(D)** GCaMP baseline (-0.009 ± 0.3, n=14) does not increase after Chrimson activation (0.04 ± 0.3, n=44). Non-parametric paired Wilcoxon test, N.S., Not Significant, p = 0.305.

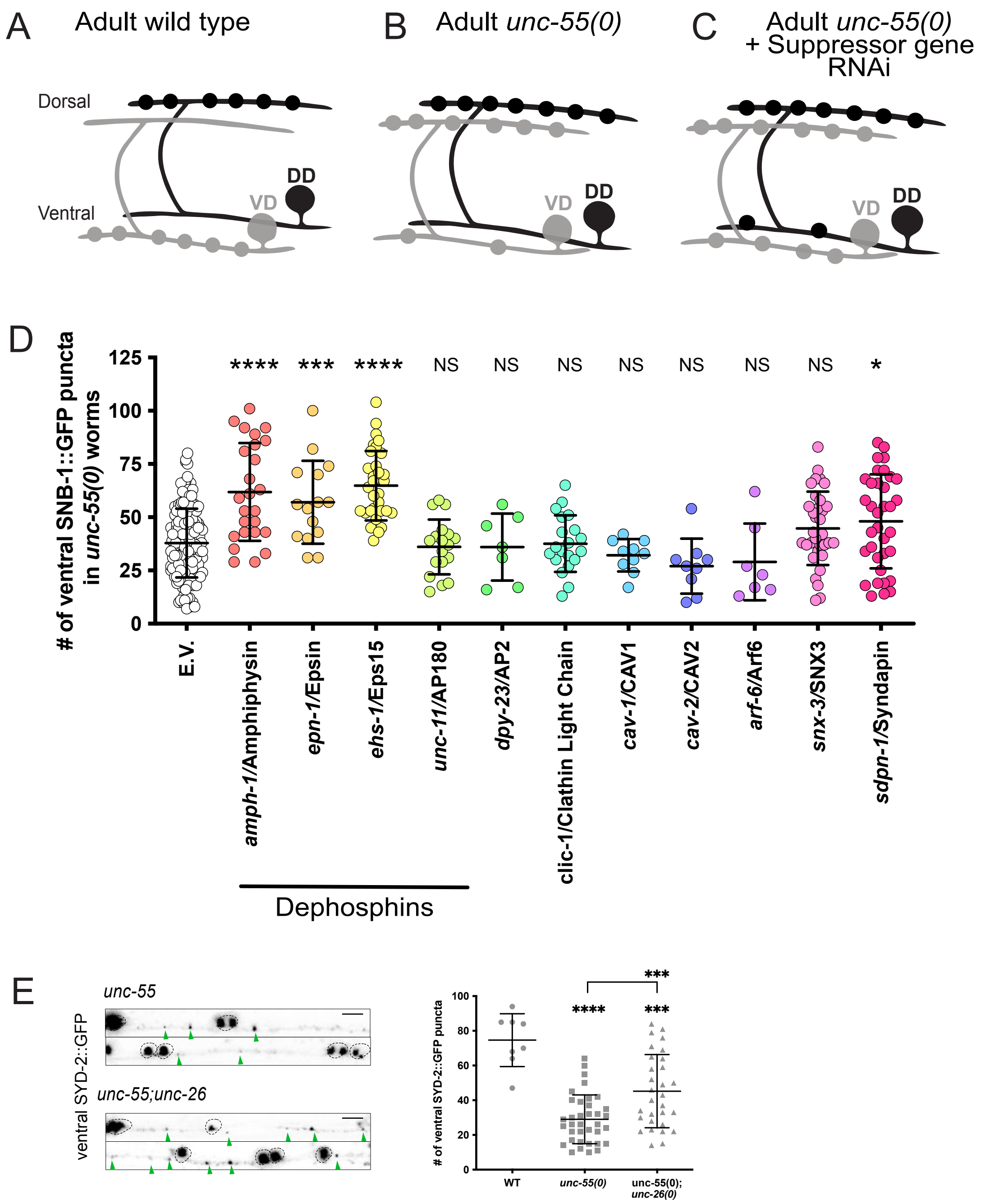

A
Adult wild type
Dorsal
Ventral
VD
DD
B
Adult unc-55(0)
VD
DD
C
Adult unc-55(0)
+ Suppressor gene
RNAi
VD
DD
D
### of ventral SNB-1::GFP puncta
in unc-55(0) worms
125
100
75
50
25
0
**** *** **** NS NS NS NS NS NS NS *
E.V.
amph-1/Amphiphysin
epn-1/Epsin
ehs-1/Eps15
unc-11/AP180
dpy-23/AP2
clic-1/Clathin Light Chain
cav-1/CAV1
cav-2/CAV2
arf-6/Arf6
snx-3/SNX3
sdpn-1/Syndapin
Dephosphins
E
unc-55
unc-55;unc-26
ventral SYD-2::GFP
### of ventral SYD-2::GFP puncta
100
80
60
40
20
0
****
***
***
WT
unc-55(0)
unc-55(0);
unc-26(0)

**Figure 4 - Supplement. Dephosphins are required for efficient removal of presynaptic SNB-1/Synaptobrevin in remodeling GABAergic neurons.**

**A-C.** Presynaptic boutons for DD (black) and VD (gray) neurons in adults. (**A**) In the wild type, DD boutons localize to the dorsal nerve cord and VD boutons are positioned on the ventral side. (**B**) In *unc-55* mutant animals, VD neurons ectopically remodel, eliminating ventral synapses and locating boutons on the dorsal side. (**C**) RNAi knockdown of genes required for the Unc-55 phenotype (i.e., “Suppressor genes”), results in the retention of ventral GABAergic synapses.

**D.** Number of ventral presynaptic boutons (SNB-1::GFP) in *unc-55* mutant animals treated with control RNAi (37.9 ± 16.2, n=153) or RNAi knockdown of dephosphins, *amph-1* (61.8 ± 22.9, n=25), *epn-1* (57.0± 19.5, n=15) and *ehs-1* (64.8 ± 16.3, n=33); regulators of clathrin-mediated endocytosis, *unc-11* (36.1 ± 12.9, n=18), *dpy-23* (36.0 ± 15.7, n=7), *clic-1* (37.6 ± 13.2, n=20); caveolins: *cav-1* (32.1 ± 7.6, n=10) and *cav-2* (27.0 ± 12.9, n=9) and of *arf-6* (29.0 ± 18.1, n=7), retromer component *snx-3* (44.8 ± 17.3, n=34) and *sdpn-1* (48.1 ± 22.1, n=35). One-Way ANOVA with Dunnett’s multiple comparison test. **** <0.0001, *** <0.001, * <0.05, NS is Not Significant.

**E.** Ventral SYD-2::GFP puncta are retained in adult *unc-55* mutants animals (29 ± 14.1, n=36) when dephosphin Synaptojanin/*unc-26* is mutated (45.2 ± 21.1, n=30). Wild-type animals (74.6 ± 15.2, n=8) have more ventral SYD-2::GFP puncta than *unc-55; unc-26.* One-Way ANOVA with Dunnett’s multiple comparison test. **** <0.0001 and *** <0.001. Scale bar = 5 μm.

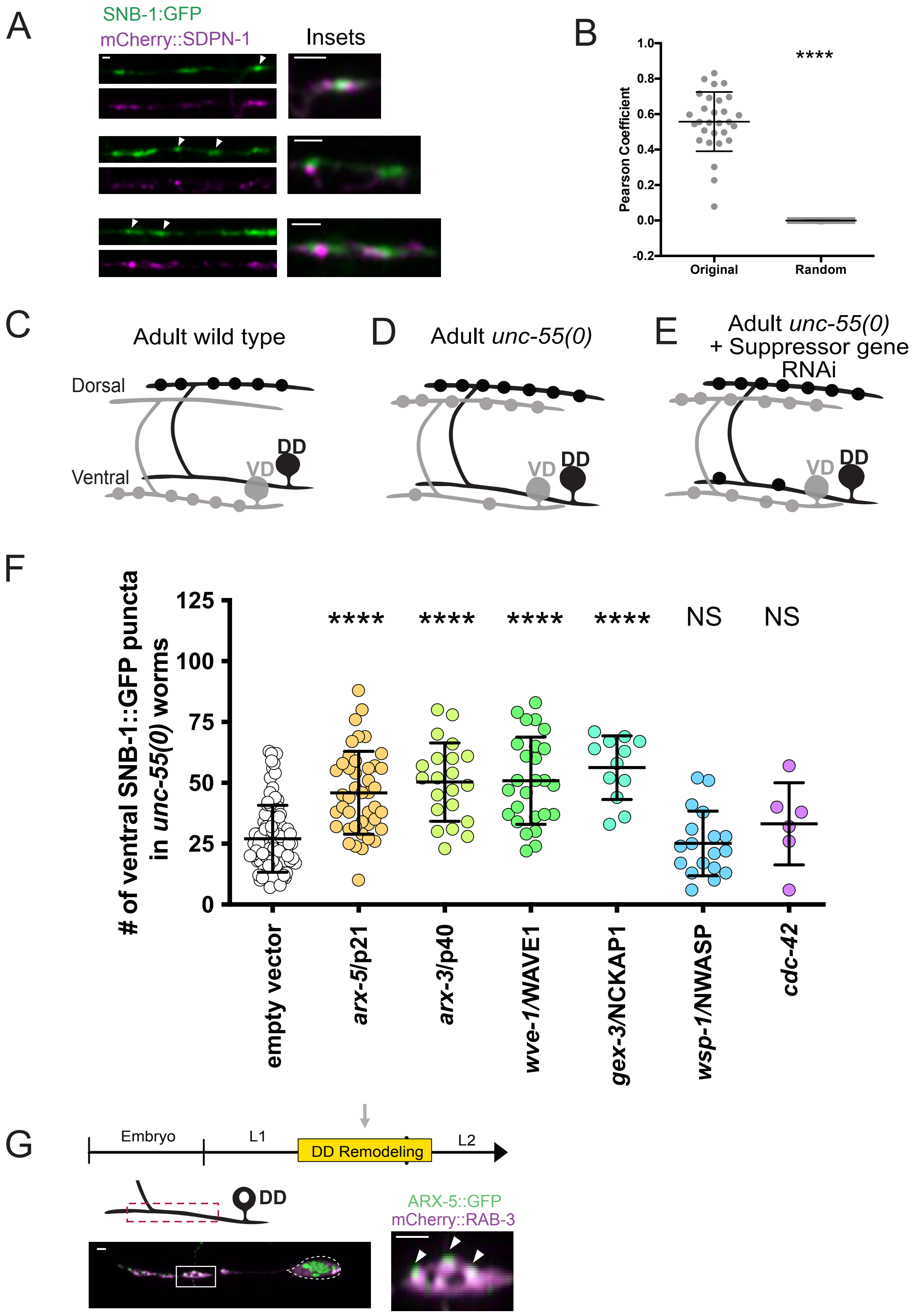

A
SNB-1:GFP
mCherry::SDPN-1
Insets
B
Pearson Coefficient
****
Original
Random
C
Adult wild type
Dorsal
Ventral
DD
VD
D
Adult unc-55(0)
E
Adult unc-55(0)
+ Suppressor gene
RNAi
F
### of ventral SNB-1::GFP puncta
in unc-55(0) worms
**** **** **** **** NS NS
empty vector
arx-5/p21
arx-3/p40
wve-1/WAVE1
gex-3/NCKAP1
wsp-1/NWASP
cdc-42
G
Embryo
L1
DD Remodeling
L2
DD
ARX-5::GFP
mCherry::RAB-3

**Figure 5 – Supplement. Syndapin and branched-actin localize to GABAergic neuron synapse and promote synaptic remodeling.**

**A.** Airyscan imaging detects SDPN-1::mCherry (magenta) localized with presynaptic clusters labeled with SNB-1::GFP (green) in remodeling L2 stage DD neurons. Insets feature representative synapses (arrowheads). Scale bar = 2 μm.

**B.** 3D Z-stacks were subjected to JACoP analysis, Pearson Coefficient between SDPN-1::mCherry and SNB-1::GFP signal is 0.56 ± 0.2 for original images and 0 ± 0.0004 for a randomized array of pixels in 3D. Mann Whitney test, *** is $p < 0.0001$. N = 29 boutons.

**C-E.** Presynaptic boutons for DD (black) and VD (gray) neurons in adult animals. (**A**) In wild-type animals, DD boutons localize to the dorsal nerve cord and VD boutons are positioned on the ventral side. (**B**) In *unc-55* mutant animals, VD neurons ectopically remodel, eliminating ventral synapses and locating boutons on the dorsal side. (**C**) RNAi knockdown of genes required for the Unc-55 phenotype (i.e., "Suppressor genes"), results in the retention of ventral GABAergic synapses.

**F.** Number of ventral synapses in *unc-55* mutant animals subjected to control RNAi (27.0 ± 13.7, n=103) or RNAi knockdown of Arp2/3 subunits, *arx-5* (45.9 ± 16.9, n=43) and *arx-3* (50.3 ± 16.1, n=21). Knockdown of subunits of the Wave Regulatory Complex, *wve-1* (50.9 ± 17.9, n=26) and *gex-3* (56.3 ± 13.0, n=12). Knockdown of *WASP/wsp-1* (25.1 ± 13.3, n=18) and GTPase *cdc-42* (33.2 ± 16.9, n=6). One-Way ANOVA with Dunnett's multiple comparison test. **** <0.0001, *** <0.001, NS is not significant.

**G.** Airyscan imaging of ventral DD neurites during remodeling (grey arrow) shows ARX-5::GFP (green) puncta (arrowheads) adjacent to synaptic vesicle clusters labeled with mCherry::RAB-3 (magenta) (inset). Dashed line denotes DD cell soma. Scale bar = 2 μm.

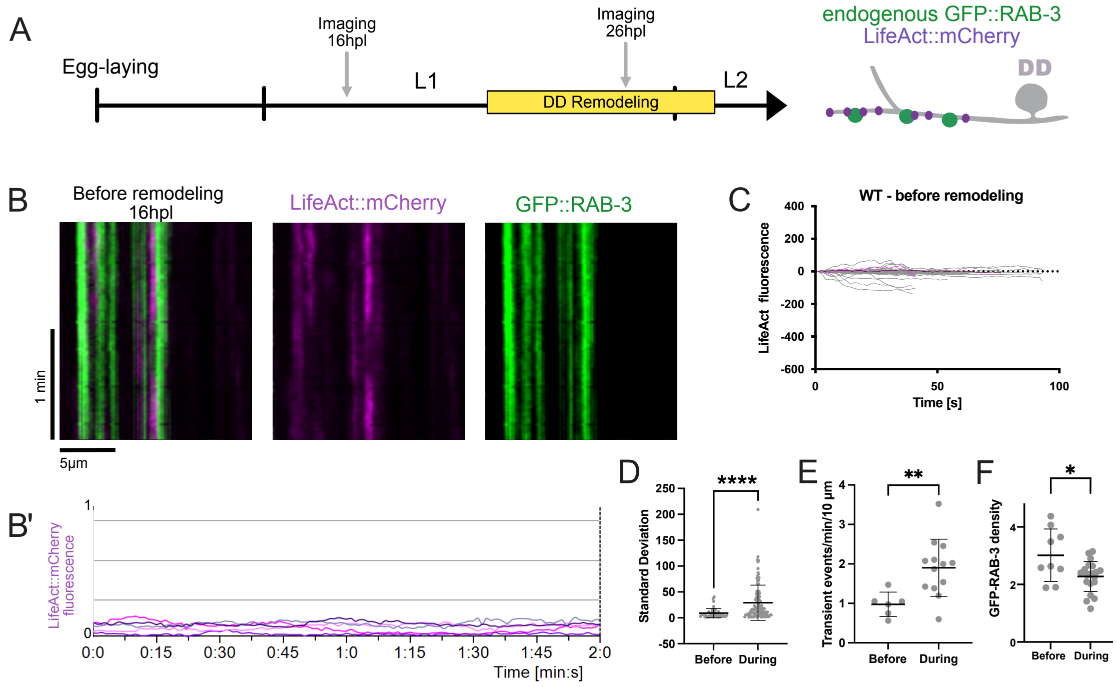
A
Egg-laying
Imaging
16hpl
Imaging
26hpl
L1
DD Remodeling
L2
endogenous GFP::RAB-3
LifeAct::mCherry
DD
B
Before remodeling
16hpl
LifeAct::mCherry
GFP::RAB-3
1 min
5µm
B'
LifeAct::mCherry
fluorescence
Time [min:s]
C
WT - before remodeling
LifeAct fluorescence
Time [s]
D
Standard Deviation
****
Before
During
E
Transient events/min/10 µm
**
Before
During
F
GFP-RAB-3 density
*
Before
During

**Figure 6 – Supplement. Upregulation of actin dynamics and GFP::RAB-3 mobility during DD remodeling.**

**A.** Ventral regions of DD neurons were imaged in L1 larvae before (16hpl) and during remodeling (26hpl) (gray arrows) to monitor actin dynamics with LifeAct::mCherry (magenta) and synaptic vesicles with endogenous GFP::RAB-3 (green).

**B.** Before remodeling (16 hpl), kymographs from live-imaging videos at 1fps (frame per second) show synaptic LifeAct::mCherry associated with stable GFP::RAB-3 puncta. B' Line scans from kymograph show stable LifeAct::mCherry fluorescence, n = 5 boutons. The Y-axis is normalized to the maximum value of LifeAct:mCherry fluorescence in line scans shown in Figure 6 C'.

**C.** Synaptic LifeAct::mCherry fluorescence is stable before remodeling (N = 6 videos).

**D.** The standard deviation of presynaptic LifeAct::mCherry signals before remodeling (9.04 ± 9.0, n=43 boutons from 6 videos) is significantly elevated during remodeling (29.1 ± 34.0, n=102 boutons from 14 videos).

**E.** More transient GFP::RAB-3 events were detected during DD remodeling (1.86 ± 0.8, n = 13 videos) than before the remodeling period (0.93 ± 0.7, n = 6 videos).

**F.** Density of GFP::RAB-3 puncta (puncta/10 microns) is higher before remodeling (3.01 ± 0.9, n = 9 snapshots) than during the DD remodeling window (L2 larval stage) (2.28 ± 0.5, n = 21 snapshots). T-test, * is $p<0.05$ and ** is $p<0.01$.

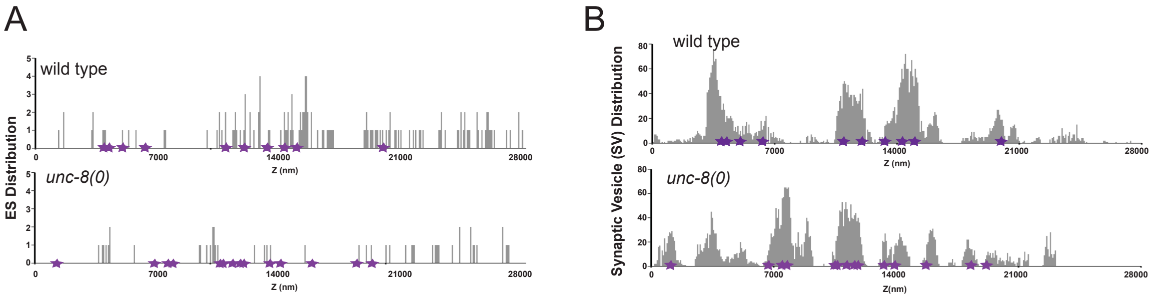


C

| Strain | Neurite Length (um) | # Dense Projection (DP) | Avg. Vol. of DP ± SEM (nm³) | Endocytic Structures (ES) | Avg. Vol. of ES ± SEM (nm³) | Mitochondria |
|---|---|---|---|---|---|---|
| **wild type** | 27.72 | 10 | $28.9 \pm 8.5 \times 10^4$ | 133 | $22.5 \pm 3.4 \times 10^4$** | 3 |
| ***unc-8(0)*** | 24.36 | 14 | $42.7 \pm 14.6 \times 10^4$ | 55 | $41.2 \pm 5.2 \times 10^4$** | 5 |

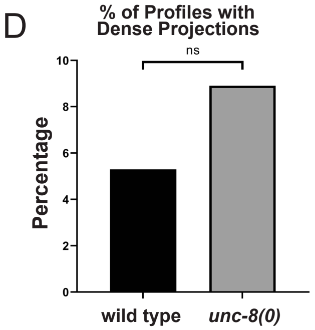


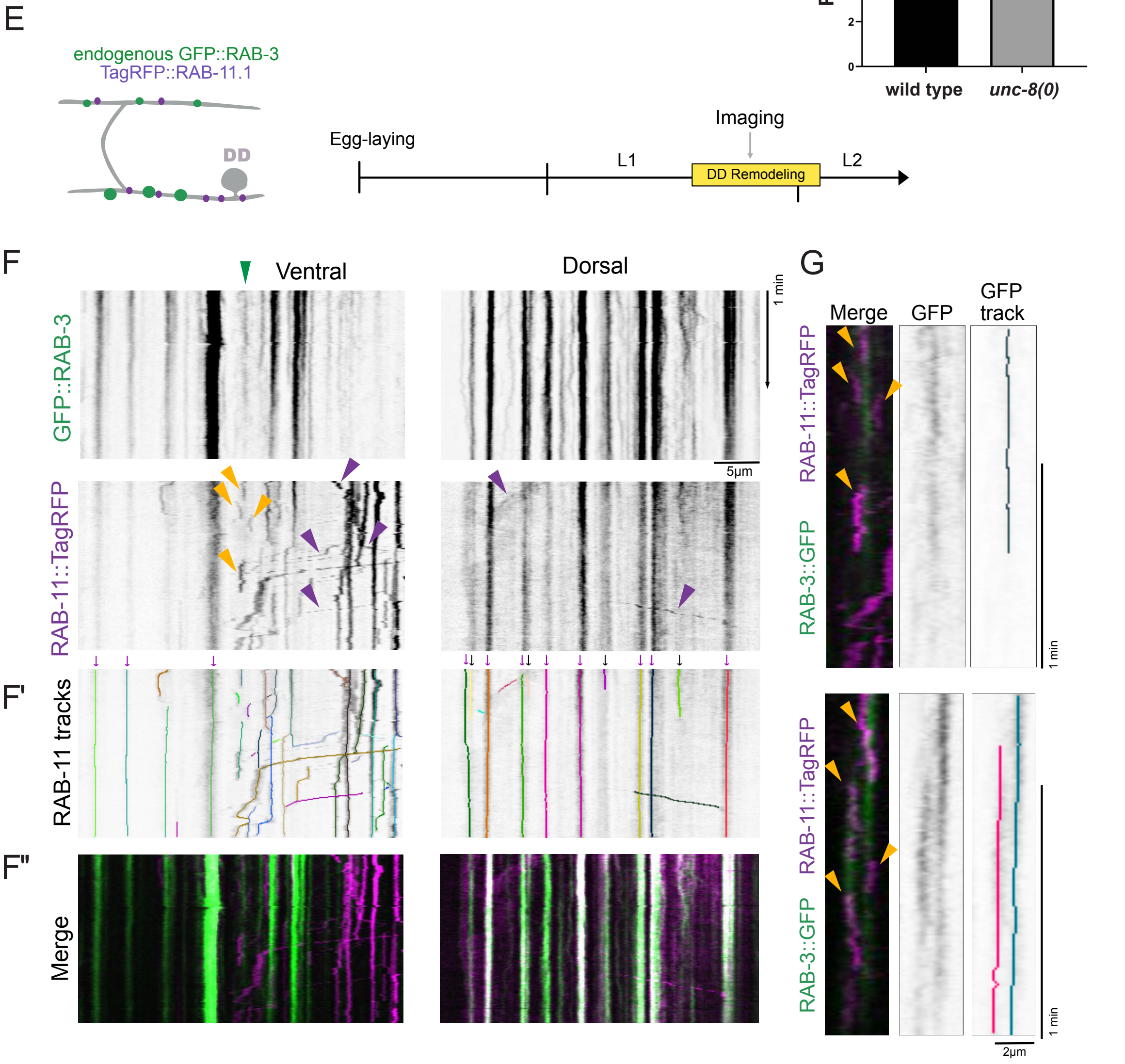

**Figure 7 – Supplement. RAB-11 endosome trafficking in remodeling DD neurites.**

**A.** Locations of endocytic structures along the lengths of the reconstructed wild-type and *unc-8(0)* DD2 neurons. Purple stars denote locations of dense projections. Z=0 is anterior most TEM section of the neuron. nm = nanometer.

**B.** Distribution of Synaptic Vesicles (SVs) (grey) along the lengths of reconstructed wild-type and *unc-8(0)* DD2 neurons. Purple stars denote locations of dense projections. Z=0 is anterior most TEM section of the neuron. nm = nanometer.

**C.** Quantifications from reconstructed wild-type and *unc-8(0)* DD2 neurons. Statistical significance was calculated with an unpaired student t-test (** p=0.0061).

**D.** Percentage of TEM profiles (sections) containing dense projections in reconstructed DD2 neurons. Statistical significance was calculated with Fischer's Exact Test, ns = not significant.

**E.** (Left) Synaptic vesicles (GFP::RAB-3) and recycling endosomes (TagRFP::RAB-11.1) were imaged (Right) in ventral cord DD neurites during the remodeling window (arrow).

**F.** Kymographs from ventral (Left) and dorsal (Right) neurites of remodeling DD neurons. (Bottom) Dynamic TagRFP::RAB-11 particles interact with (Top) GFP::RAB-3 puncta. Kymographs reveal mobile RAB-11 particles (purple arrowheads) and RAB-11 (yellow arrowheads) that associate with transient GFP::RAB-3 on the ventral side (green arrowhead).

**F'** RAB-11 particles in dorsal and ventral cords identified with Kymobutler (See Methods)

**F''**. Merged images of RAB-11 (Magenta) and RAB-3 (green) particles

**G.** (Left) Representative kymographs showing dynamic association of RAB-11 (yellow arrowheads) with transient GFP::RAB-3 puncta (green) in ventral DD neurites. (Right) Kymographs of transient GFP::RAB-3 with Kymobutler tracks.

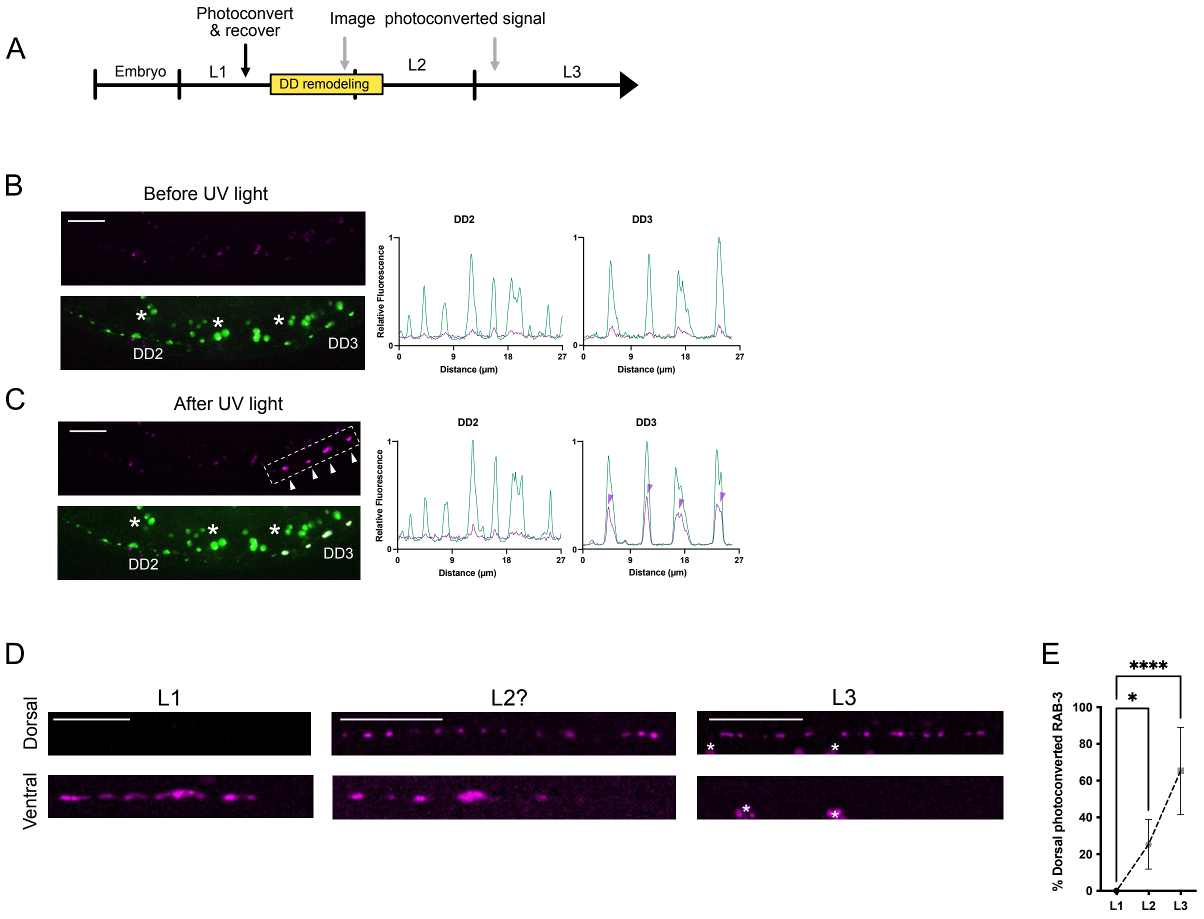
A
Photoconvert & recover
Image photoconverted signal
Embryo
L1
DD remodeling
L2
L3
B
Before UV light
DD2
DD3
Relative Fluorescence
Distance (μm)
C
After UV light
DD2
DD3
Relative Fluorescence
Distance (μm)
D
L1
L2?
L3
Dorsal
Ventral
E
% Dorsal photoconverted RAB-3
****
*
L1
L2
L3

**Figure 8 – Supplement. Progressive recycling of RAB-3 from old to new synapses in remodeling DD neurons.**

**A.** Experimental set up: Dendra-2::RAB-3 was photoconverted in wild type animals (black arrow) and worms were recovered for imaging either during remodeling (L2) (gray arrow) or after remodeling (L3) (gray arrow).

**B.** (Left) Before UV-irradiation, only green fluorescence is detected in the ventral cord. (Right) Line tracings of ventral DD2 and DD3 neurites. Asterisks mark auto fluorescent granules in the intestine.

**C.** (Left) After UV-irradiation of the ventral DD2 neurite (dashed box) green and photoconverted Dendra-RAB-3 puncta are detected in the ventral cord. (Right) Line tracings convert photoconversion of Dendra-RAB-3 in ventral DD2 but not DD3 neurites. Asterisks mark auto fluorescent granules in the intestine.

**D.** Representative images of photoconverted Dendra-2::RAB-3 puncta in dorsal and ventral nerve cords immediately after photoconversion (L1), during remodeling (late L2) and after remodeling (L3). Asterisks denote autofluorescence.

**E.** Progressive accumulation of photoconverted Dendra-2::RAB-3 on the dorsal side : L1 ($0 \pm 0$, $n = 17$), at L2 ($25.3 \pm 13.5$ %, $n = 6$) and at L3 ($65.3 \pm 23.7$ %, $n = 14$). Kruskall-Wallis test with Dunn's multiple comparison test. * $p = 0.019$ and *** $p < 0.0001$. Scale bars = 10 μm.
