## Supplementary material for "The Epithelial Na^+^ Channel UNC-8 promotes an endocytic mechanism that recycles presynaptic components from old to new boutons in remodeling neurons": Video Legends

### VIDEOS

**Video S1. Spontaneous calcium dynamics in the ventral anterior neurite of remodeling wild-type DD neuron**

(Top) GCaMP6s expressed in DD neurons and recorded at 10 fps in the ventral DD neurite of a wild-type animal during remodeling (26 hpl). ROIs were defined at the remodeling bouton (yellow) and at the neurite (magenta) adjacent to the DD cell body. (Bottom) GCaMP fluorescent intensity displayed across time show bouton-specific $Ca^{++}$ transients. Scale bar = 5 μm.

**Video S2. Spontaneous calcium dynamics in the ventral anterior neurite of remodeling *unc-8* mutant DD neuron**

(Top) GCaMP6s expressed in DD neurons and recorded at 10 fps in the ventral DD neurite of an *unc-8* mutant animal during remodeling (26 hpl). ROIs were defined at the remodeling bouton (yellow) and at the neurite (magenta) adjacent to the DD cell body. (Bottom) GCaMP fluorescent intensity displayed across time show bouton-specific $Ca^{++}$ transients. Scale bar = 5 μm.

**Video S3. Evoked calcium transients in the ventral anterior neurite of remodeling wild-type and *unc-8* DD neurons.**

Evoked calcium transients upon activation of the presynaptic VA neurons (*Punc-4::*Chrimson) during remodeling (26 hpl) in wild-type (top) and *unc-8* mutant (bottom) animals. Magenta dot at the left upper corner marks the video frame immediately after optogenetic activation. Arrows point to the remodeling bouton used for quantitative analysis (Figure 2). Scale bar = 2 μm. Recorded at 10 fps.

**Video S4. Actin and RAB-3 dynamics before remodeling in a wild-type animal**

Video (1 fps) of actin dynamics (magenta) in presynaptic regions marked with RAB-3 (green) in DD neuron before remodeling (16 hpl). Scale bar = 1 μm.

**Video S5. Actin and RAB-3 dynamics during remodeling in a wild-type animal**

Video (1 fps) of actin dynamics (magenta) in presynaptic regions marked with RAB-3 (green) in DD neuron during remodeling period (26 hpl). Note strong actin dynamics and mobile RAB-3 puncta in inter-synaptic regions. Scale bar = 2 μm.

**Video S6. Actin polymerization coincides with RAB-3 disappearance**

(Top) Video (1 fps) of dynamic GFP::RAB-3 punctum (arrow) that dissipates with burst of actin (LifeAct) signal (magenta). (Bottom) Quantification of GFP:RAB-3 and LifeAct::mCherry fluorescence at transient GFP::RAB-3 punctum (arrow)

**Video S7. Actin and RAB-3 dynamics during remodeling in an *unc-8* mutant**

Video (1 fps) of actin dynamics (magenta) at presynaptic regions, labeled with RAB-3 (green) during the remodeling window (26 hpl) of an *unc-8* mutant. Scale bar = 2 μm.

**Video S8. EM reconstruction of ventral DD2 neurite during remodeling**
3D rendering of EM sections  control anterior ventral DD2 neurite during remodeling (see Figure 7B).

**Video S9. RAB-11 dynamics during DD remodeling**

TagRFP::RAB-11  puncta (magenta) travel along the DD commissure (arrows) during DD remodeling (26 hpl).
